# An Explainable AI Framework for Identifying Universal Aging Signatures in Cell Embeddings

**DOI:** 10.1101/2025.11.07.687286

**Authors:** Wei Qiu, Chris Arian, Ethan Weinberger, Soo R. Yun, Alexander R. Mendenhall, Jessica E. Young, Maria Brbić, Su-In Lee

## Abstract

Aging is a complex biological process marked by progressive physiological decline and increased disease vulnerability. Single-cell RNA sequencing offers unprecedented resolution for studying aging, yet isolating aging-related signatures remains challenging because gene expression is primarily shaped by other factors such as cell type, tissue, and sex. We present **ACE (Aging Cell Embeddings)**, an explainable deep generative framework that disentangles aging-related gene expression changes from background biological variation. ACE employs two latent representations: one capturing aging-related signatures and another representing non-aging-related variation in the data. Through explainable AI, ACE identifies key genes and pathways associated with aging amid dominant non-aging-related variations. Applied to large-scale mouse, fly, and human datasets, ACE uncovers aging signatures both within specific tissue-cell-type contexts and across all tissues and cell types, enabling accurate prediction of biological age. Moreover, ACE identifies aging genes conserved across species, highlighting its ability to reveal shared biological mechanisms of aging. Experimental RNAi knockdowns in *C. elegans* validate ACE’s findings, confirming its ability to prioritize novel aging genes affecting lifespan. ACE reveals key pathways involved in proteostasis, immune regulation, and extracellular matrix remodeling, and identifies *Uba52* through the cross-species model as an important aging gene, whose knockdown in *C. elegans* significantly shortens lifespan. By providing interpretable and generalizable aging embeddings, ACE establishes a foundation for cross-species single-cell aging studies and translational geroscience.

## Introduction

Aging is a complex, multifactorial process characterized by the progressive loss of physiological integrity, which leads to functional decline and increased vulnerability to disease [1]. It is the primary risk factor for a wide range of chronic diseases, including cancer, diabetes, cardiovascular disorders, and neurodegenerative conditions [2, 3]. Unraveling the biological mechanisms that drive aging is essential for identifying therapeutic targets and biomarkers that can promote healthy longevity and delay aging-related diseases [4, 5]. Although numerous studies have uncovered molecular and cellular changes associated with aging [6, 7], identifying the key drivers remains one of the field’s most pressing challenges [8].

The advent of single-cell transcriptomic technologies has enabled unprecedented resolution in exploring the aging process. By capturing gene expression profiles at the individual cell level, single-cell RNA sequencing (scRNA-seq) facilitates the discovery of cell-type-specific and context-dependent aging signatures [9, 10, 11]. However, accurately characterizing aging signatures from scRNA-seq data remains challenging due to the other non-aging-related sources of variation such as tissue, cell type, sex, and batch effects. These background factors often obscure the more subtle transcriptomic changes associated with aging, hindering efforts to isolate true aging-related signatures. To address this complexity, previous studies have developed separate linear models for different tissues and cell types [12, 13, 14], an approach that requires training many tissue- and cell-specific models and risks overlooking globally conserved aging genes.

Recent single-cell representation learning methods, including probabilistic latent variable models and contrastive approaches, capture broad cellular heterogeneity but generally do not disentangle subtle aging-related signatures from other dominant biological factors [15, 16, 17, 18, 19, 20, 21]. These frameworks often rely on a single latent space or on binary case–control contrasts, which do not reflect the continuous nature of aging. In recent disentanglement frameworks [22], all cells belonging to the same category, such as a specific cell type or condition, share a single embedding, making it difficult to capture gradual, continuous processes such as aging. Together, these limitations motivate an approach that explicitly models aging-related and background variation.

To address these challenges, we introduce ACE (Aging Cell Embeddings), a representation learning framework designed to disentangle aging-related signatures from background biological variation in scRNA-seq data (Figure 1a). ACE models gene expression using two distinct sets of latent variables: *age variables*, which capture variations related to age, and *background variables*, which capture other non-aging-related signatures (Figure 1b). The framework supports both global (*i.e*., shared across tissues and cell types) and local (*i.e*., tissue-cell-type-specific) aging analyses, enabling the identification of both shared and context-specific aging trajectories.

**Figure 1.**
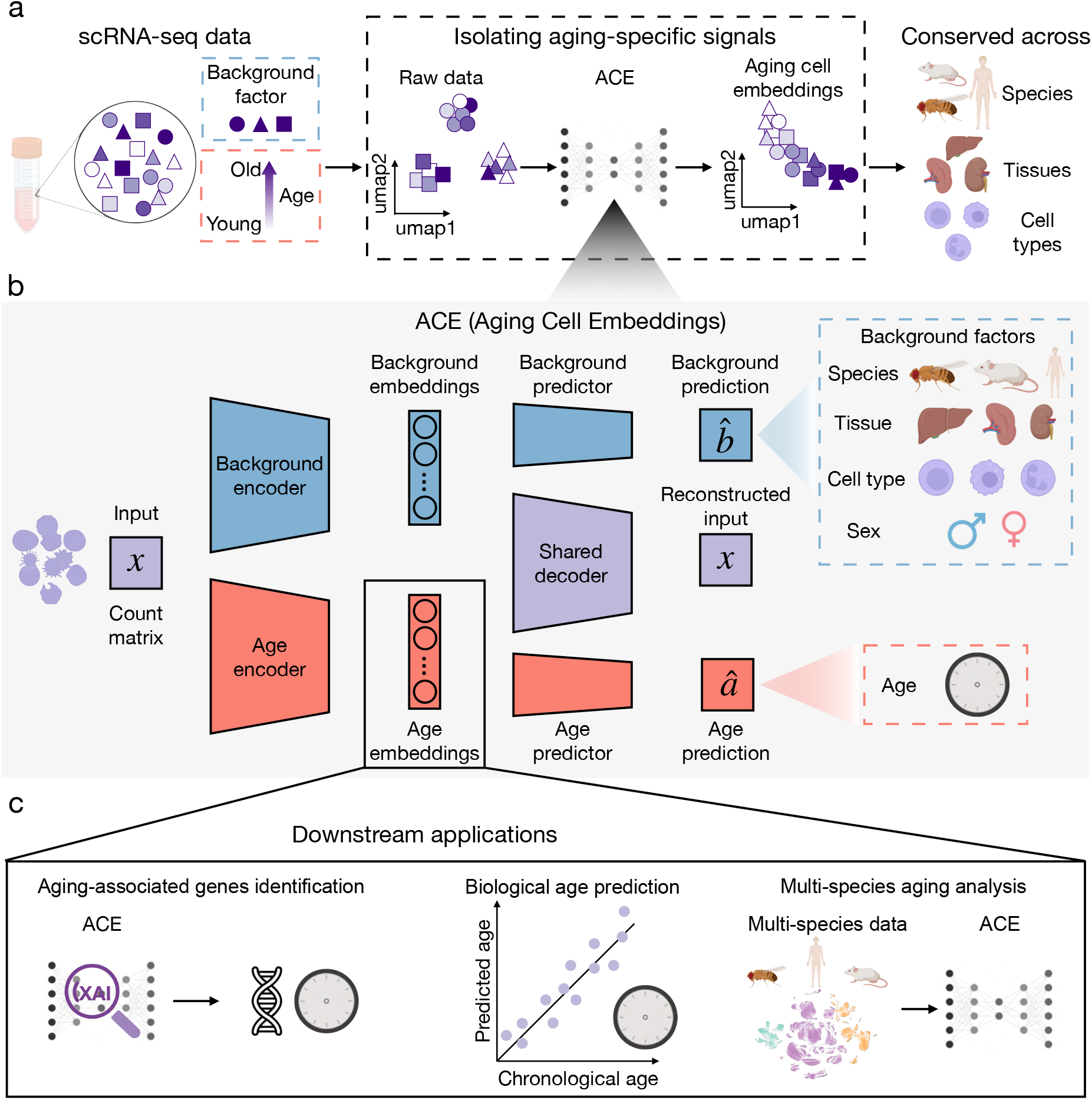
Overview of the ACE framework for disentangling aging signatures in scRNA-seq data. **a.** ACE is designed to isolate aging-associated gene expression signatures from raw scRNA-seq data, which are often confounded by other biological variation such as species, tissue, cell type, and sex (*i.e*., background factors). By disentangling aging signatures from the background factors, ACE learns aging-associated cell embeddings that represent cellular aging independently of context. ACE is applicable to large-scale datasets that include multiple cell types, tissues, and species, enabling the characterization of aging-associated changes across different biological axes. This design allows for systematic and scalable aging analysis in heterogeneous single-cell datasets. **b**. ACE consists of two parallel encoders: an age encoder that extracts latent variables capturing aging-associated signatures and a background encoder that extracts latent variables capturing dominant background factors. These variables are used by corresponding predictors to reconstruct background labels and chronological age, respectively. A shared decoder reconstructs the input gene expression matrix, encouraging the two sets of latent variables to capture complementary information. The explicit separation of aging and background signatures allows for robust modeling of aging trajectories across heterogenous single-cell datasets. **c**. The disentangled aging representations learned by ACE enable a variety of downstream applications, such as (1) identification of global (*i.e*., shared across tissues and cell types) and local (*i.e*., tissue-cell-type-specific) aging-associated genes and pathways using explainable AI (XAI) methods; (2) prediction of biological age from gene expression profiles; and (3) analysis of aging patterns conserved across different species by aligning their aging trajectories.

We apply ACE to large-scale single-cell aging atlases, including datasets from mouse [9], fly [10], and human [23]. Our results show that ACE effectively isolates aging-related signatures and enables several key applications (Figure 1c): (1) identifying global and local aging genes using explainable artificial intelligence (XAI) methods; (2) predicting biological age at both the cell and subject levels across diverse contexts; and (3) analyzing conserved aging signatures across species, including mouse, fly, and human, by aligning aging trajectories and identifying conserved aging genes. We further validate ACE’s robustness through cross-cell-type generalization, highlighting its utility in uncovering context-independent aging mechanisms. Finally, we experimentally confirmed ACE’s findings through RNAi knockdown experiments in *C. elegans*, where multiple prioritized genes significantly impacted lifespan. Notably, ACE identified *Uba52* as an aging-associated gene across species, and its role was experimentally validated, demonstrating ACE’s ability to uncover biologically conserved aging mechanisms. The implementation of the ACE model is publicly available online.^1^

## Results

### ACE effectively disentangles aging signatures from single-cell RNA-seq data

We designed ACE (Aging Cell Embeddings), an explainable variational autoencoder-based model, to isolate aging-related gene expression variations in single-cell RNA sequencing datasets (Figure 1a). scRNA-seq datasets are inherently complex, with gene expression signatures reflecting diverse biological factors including species, tissue, cell type, sex, and technical artifacts. ACE is a representation learning framework [24] that explicitly models and disentangles aging-related variations from other non-aging-related factors in scRNA-seq data. It enables the extraction of conserved aging signatures from heterogeneous biological contexts.

The architecture of ACE (Figure 1b; Methods) comprises two parallel encoder networks. The age encoder extracts *age embeddings* that specifically capture aging-related signatures. Simultaneously, the background encoder captures embeddings that represent non–aging-related variation, such as species, tissue, cell type, and sex (*i.e*., background factors). To ensure effective separation of these embeddings, ACE incorporates prediction networks: age embeddings are used to predict the subjects’ chronological age, while background embeddings predict background factors. The two sets of embeddings are constrained to remain independent using the Hilbert-Schmidt Independence Criterion (HSIC) [25, 26, 27], ensuring that the learned biological signatures remain distinct. To reconstruct gene expression while accounting for technical variation, ACE includes a decoder that models the observed RNA measurements using distributions conditioned on the background and age embeddings. These distributions account for technical factors such as sequencing depth, batch effects, and dropouts by following the probabilistic modeling approach used in scVI [15].

The age variables learned by ACE enable a variety of downstream applications essential to aging research (Figure 1c). Specifically, the age embeddings support: (1) identification of global (*i.e*., shared across tissues and cell types) and local (*i.e*., tissue-cell-type-specific) aging genes using explainable AI methods, (2) accurate estimation of biological age, defined as the prediction of aging based on gene-expression profiles, at both the cell and subject levels, and (3) analysis of conserved aging signatures across different species by aligning their aging trajectories. These applications underscore the model’s utility in systematically disentangling aging signatures from complex scRNA-seq data.

### ACE effectively captures global aging trajectories

Aging manifests through both global and local molecular programs: some transcriptional signatures are shared across diverse tissues and cell types, whereas others are restricted to particular cellular contexts. ACE supports global aging analysis by explicitly modeling aging-related variation that is consistent across tissues and cell types (Figure 2a). For this analysis, ACE was trained separately on each species (*e.g*., mouse and fly), with tissue, cell type, and sex as background factors, allowing the age embeddings to capture variation specifically associated with aging while controlling for other biological sources of heterogeneity. We evaluated ACE’s ability to disentangle global aging signatures using two cross-tissue, multi-cell-type aging atlases — the Tabula Muris Senis (TMS) mouse dataset [9] and the Aging Fly Cell Atlas [10]. We applied Expected Gradients [28] to the global ACE model to identify global aging genes that contribute significantly to aging across tissues and cell types.

**Figure 2.**
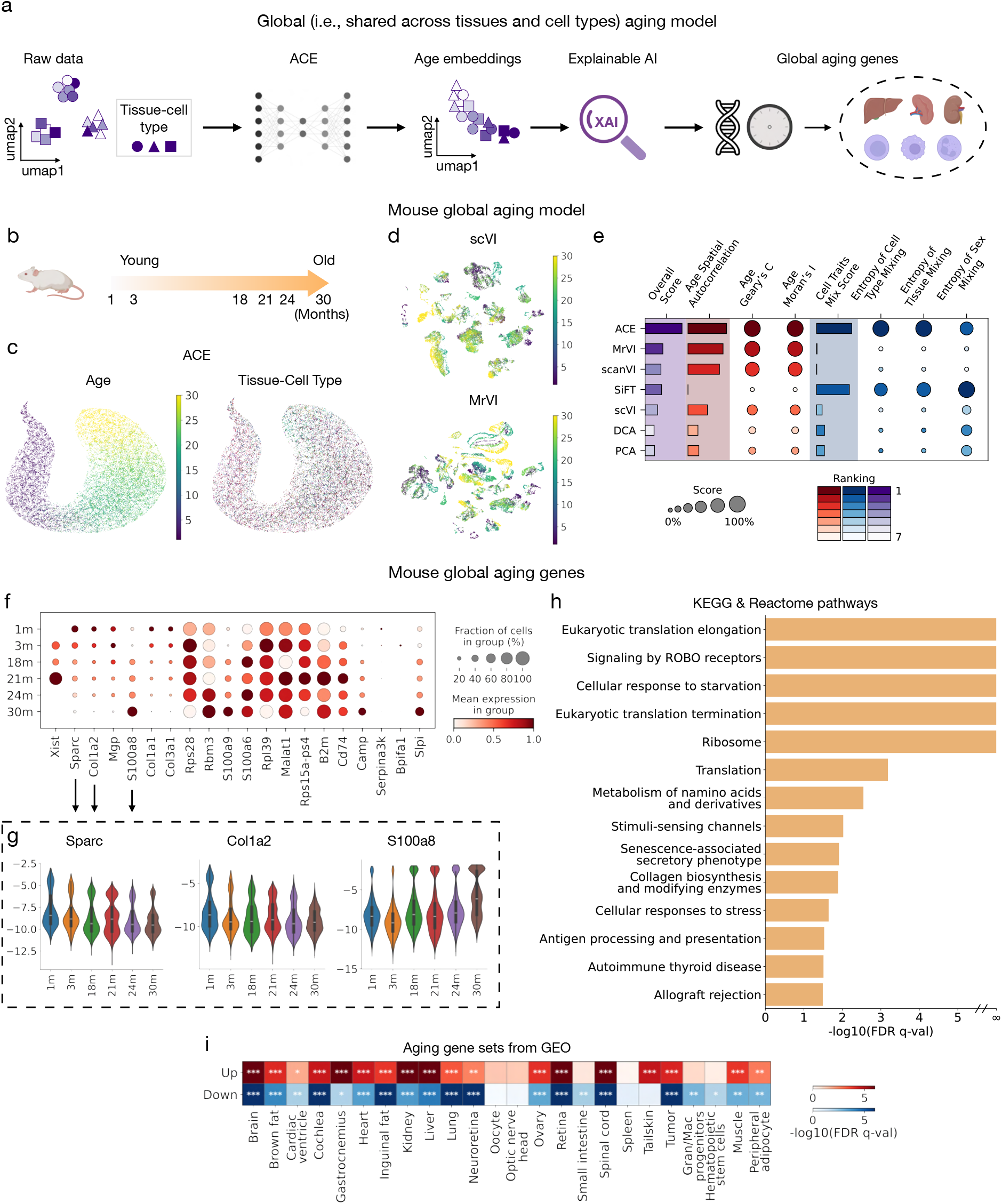
ACE enables global (*i.e*., shared across tissues and cell types) modeling of aging trajectories. **a.** Overview of the global aging model pipeline. ACE is applied to scRNA-seq data to learn aging-related cell embeddings shared across tissues and cell types. XAI techniques are then used to identify aging-associated genes across tissues and cell types. **b**. Visualization of aging timepoints available in the TMS Droplet mouse dataset, from 1 to 30 months, capturing the life span of a mouse. **c**. UMAPs of the age embeddings learned by ACE, colored by age (left) and tissue-cell-type identity (right). The age embeddings capture aging-related signatures while being disentangled from background factors. **d**. UMAPs of latent embeddings learned by alternative methods: scVI (top) and MrVI (bottom), colored by age. **e**. Quantitative comparison of ACE and baseline methods (MrVI, scanVI, SiFT, scVI, DCA, and PCA) across multiple evaluation metrics. Circles represent individual metric scores, with circle size indicating the normalized score (scaled between 0 and 1) and color reflecting the relative ranking. For consistency, all metrics are scaled such that higher values indicate better performance. Bars represent composite scores: Age Spatial Autocorrelation is the average of Geary’s C and Moran’s I; Cell Traits Mix Score is the average of cell type, tissue, and sex mixing metrics; and Overall Score is the average of Age Spatial Autocorrelation and Cell Traits Mix Score. Bar color indicates the method’s relative ranking. **f**. Dot plot showing expression patterns of top 20 global aging genes identified by the ACE global aging model in the TMS Droplet dataset. Dot size indicates the fraction of cells expressing the gene in each age group; color represents mean expression. **g**. Violin plots showing expression patterns of representative global aging genes (*Sparc, Col1a2, S100a8*). Violin plots display gene expression distributions across different age groups, aggregated over all tissues and cell types. These visualizations highlight consistent aging-associated expression changes for the selected genes. **h**. Gene set enrichment analysis of the full ranked list of global aging genes reveals KEGG and selected Reactome pathways that are significantly enriched and play key roles in aging-related biological processes. Significance was assessed at FDR q < 0.05 using the Benjamini-Hochberg correction. **i**. Gene set enrichment analysis using the “Aging Perturbations from GEO Up” and “Aging Perturbations from GEO Down” gene set collections. These databases consist of gene sets curated from GEO studies comparing aged versus young samples, capturing genes consistently upregulated or downregulated with age across various tissues and cell types. ACE-derived ranked list of global aging genes shows significant enrichment in many of these aging-associated gene sets. Red boxes indicate enrichment in *upregulated* gene sets; blue boxes indicate enrichment in *downregulated* gene sets. Color intensity reflects — log_10_(FDR q-value), with significance determined by FDR-adjusted q-values using the Benjamini-Hochberg correction. Asterisks denote significance thresholds (^∗^ *p* < 0.05, ^∗∗^ *p* < 0.01, ^∗∗∗^ *p* < 0.001). Gran/Mac progenitors represent granulocyte/macrophage progenitors.

We first trained ACE on the TMS dataset generated using droplet-based scRNA-seq, including cells from mice across different ages (1m, 3m, 18m, 21m, 24m, 30m) (Figure 2b). We focus the analysis on the 20 most commonly occurring cell types. In total, the data we use contains 135,420 cells taken from 13 different tissues across 23 subjects (Methods). The raw data’s dominant biological variations include cell type, tissue, and sex (Supplementary Figure 1). An effective global aging model should learn the age embeddings that capture aging-related variations while retaining other biological variations unrelated to aging in the background embeddings. We compared ACE with existing baseline models, including MrVI [29], scANVI [30], which utilize the age labels during training, SiFT [31], which uses the background labels during training, and scVI [15], DCA [16], and PCA, which are unsupervised single-cell representation learning methods. UMAP of ACE’s age embeddings reveals a smooth and continuous aging trajectory (Figure 2c, left), while tissue-cell-types are well mixed in the same space (Figure 2c, right), indicating that ACE effectively disentangles aging signatures from background factors and captures global aging signatures shared across tissues and cell types (see Supplementary Figure for UMAPs of both age and background embeddings colored by age and background factors).

By contrast, embeddings from baseline methods, such as scVI and MrVI, exhibited significant confounding due to tissue and cell type variations, obscuring aging trajectories (Figure 2d). A quantitative comparison further shows that ACE outperforms baseline methods across multiple metrics by a large margin, including spatial autocorrelation between the age embeddings and chronological age, as well as mixing scores for cell type, tissue, and sex (Figure 2e; Methods). Overall, ACE consistently ranks highest in capturing aging signatures while minimizing confounding from background factors.

We applied Expected Gradients to the global ACE model to identify genes contributing most strongly to aging-related variation (Methods). Among the top 20 global aging genes (Figure 2f), 16 have prior reports of aging-associated regulation: *Xist* [32], *Sparc* [33], *Col1a2* [34], *Mgp* [35], *S100a8* [36, 37], *Col1a1* [38], *Col3a1* [38], *Rps28* [39], *Rbm3* [40], *S100a9* [36], *S100a6* [41], *Malat1* [42], *Rps15a-ps4* [43] *B2m* [44], *Cd74* [45], and *Serpina3k* [46]. The remaining genes are enriched for pathways related to innate immune activation (*Camp* [47], *Bpifa1* [48], and *Slpi* [49]), suggesting plausible, but less systematically characterized, links to aging.

Several of these (*e.g*., *Sparc, Col1a2*, and *S100a8*) display clear, monotonic expression shifts across the lifespan (Figure 2g). *Sparc* functions as an immunometabolic checkpoint that promotes inflammation and interferon signaling, and its inhibition has been shown to reduce inflammation and extend health span during aging [33]. The *Col1a2* gene encodes for a component of Type I Collagen that is present in the extracellular matrix (ECM) of various tissues. ECM dynamics have been increasingly identified as important hallmarks of aging [50]. *Col1a2* has been identified as a key player in skin, bone, and cardiac health in various mouse disease models [51, 52, 53], suggesting that aging-associated changes in expression of *Col1a2* could lead to disruption of the health and homeostasis of these organ systems. *S100a8* is a pro-inflammatory gene upregulated with age in multiple tissues, where it promotes oxidative stress, inflammation, and cellular senescence, making it a potential driver and biomarker of aging [36]. The identification of these genes by ACE supports its ability to uncover biologically meaningful and well-conserved markers of aging.

We then performed gene set enrichment analysis using the full ranked list of global aging genes and identified multiple significantly enriched KEGG and Reactome pathways (Figure 2h; see Supplementary Figure for the full list of enriched pathways; Methods). These include translation-related processes (*e.g*., ribosome, eukaryotic translation elongation and termination), cellular stress responses (*e.g*., response to starvation), and immune-related pathways (*e.g*., antigen processing and presentation, senescence-associated secretory phenotype), extracellular matrix pathways (*e.g*., collagen biosynthesis and modifying enzymes) and axonal pathfinding (*e.g*., signaling by ROBO receptors). These findings underscore the relevance of proteostasis, stress adaptation, and immune regulation, cell-ECM interactions, and the nervous system in the aging process. Furthermore, the ranked gene list of global aging genes show significant enrichment across aging-associated gene sets curated from GEO, which are derived from comparisons between aged and young samples (Figure 2i). The broad enrichment across diverse tissues and cell types highlights ACE’s ability to capture conserved aging signatures shared across multiple biological contexts.

To externally validate the mouse global aging model, we applied ACE to the independent TMS FACS dataset (Methods). Similar to the Droplet data, the raw FACS data are dominated by tissue, cell type, and sex signatures, with age effects being comparatively subtle (Supplementary Figure 4). ACE effectively disentangles these factors, producing age embeddings that capture a smooth aging trajectory (Supplementary Figure 5b, Supplementary Figure 6). The global aging gene rankings from the FACS model are highly concordant with those from the Droplet model (Supplementary Figure 5c,d). Gene set enrichment analysis recapitulates key aging-associated pathways, including translation, stress response, and ROBO signaling (Supplementary Figure 5e, Supplementary Figure 7), and shows strong overlap with external GEO aging signatures (Supplementary Figure 5f), demonstrating the robustness and reproducibility of ACE across datasets.

We next evaluated ACE’s ability to extract global aging signatures on the Aging Fly Cell dataset [10], which includes cells spanning ages from 5 to 70 days (Figure 3a; Methods). The data we use contains 424,863 cells from two tissues (head and body) and 14 cell types across 30 subjects (see Supplementary Figure for the UMAPs of the raw data). Consistent with mouse data, the UMAP of ACE’s age embeddings exhibits a continuous aging trajectory that is largely independent of tissue and cell type differences (Figure 3b; see Supplementary Figure 9 for UMAPs of both age and background embeddings colored by age and background factors). ACE outperforms baseline models by best capturing aging-related variations while minimizing confounding effects from tissue, cell type, and sex (Figure 3c,d). ACE also identifies global aging genes with consistent aging-associated expression patterns (Figure 3e,f), including known aging-associated markers such as *lovit, Pzl*, and *Hsp26. lovit* is a synaptic protein that plays a critical role in transporting the neurotransmitter, histamine, in photoreceptor cells in insects and other arthropods. RNAi-mediated downregulation of *lovit* in flies resulted in disrupted visual transmission [54]. *Pzl* is expressed in specific locomotor neurons in flies, where mutant variants of *Pzl* result in disrupted locomotor function in fly larva [55]. *Hsp26*, which encodes a small heat shock protein, has been linked to increased resistance to oxidative stress and extended lifespan in flies [56]. Gene Ontology (GO) enrichment analysis of these genes highlights aging-associated pathways (Figure 3g). Similar to the mouse global model, the fly global model identifies several pathways involved in protein translation (*e.g*., Cytosolic ribosome, large ribosomal subunit, cytoplasmic translation) and synaptic function (*e.g*., Dendrite membrane, ionotropic glutamate receptor complex, and AMPA glutamate receptor complex). The agreement between these two analyses highlights the importance of proteostasis and nervous system function in aging.

**Figure 3.**
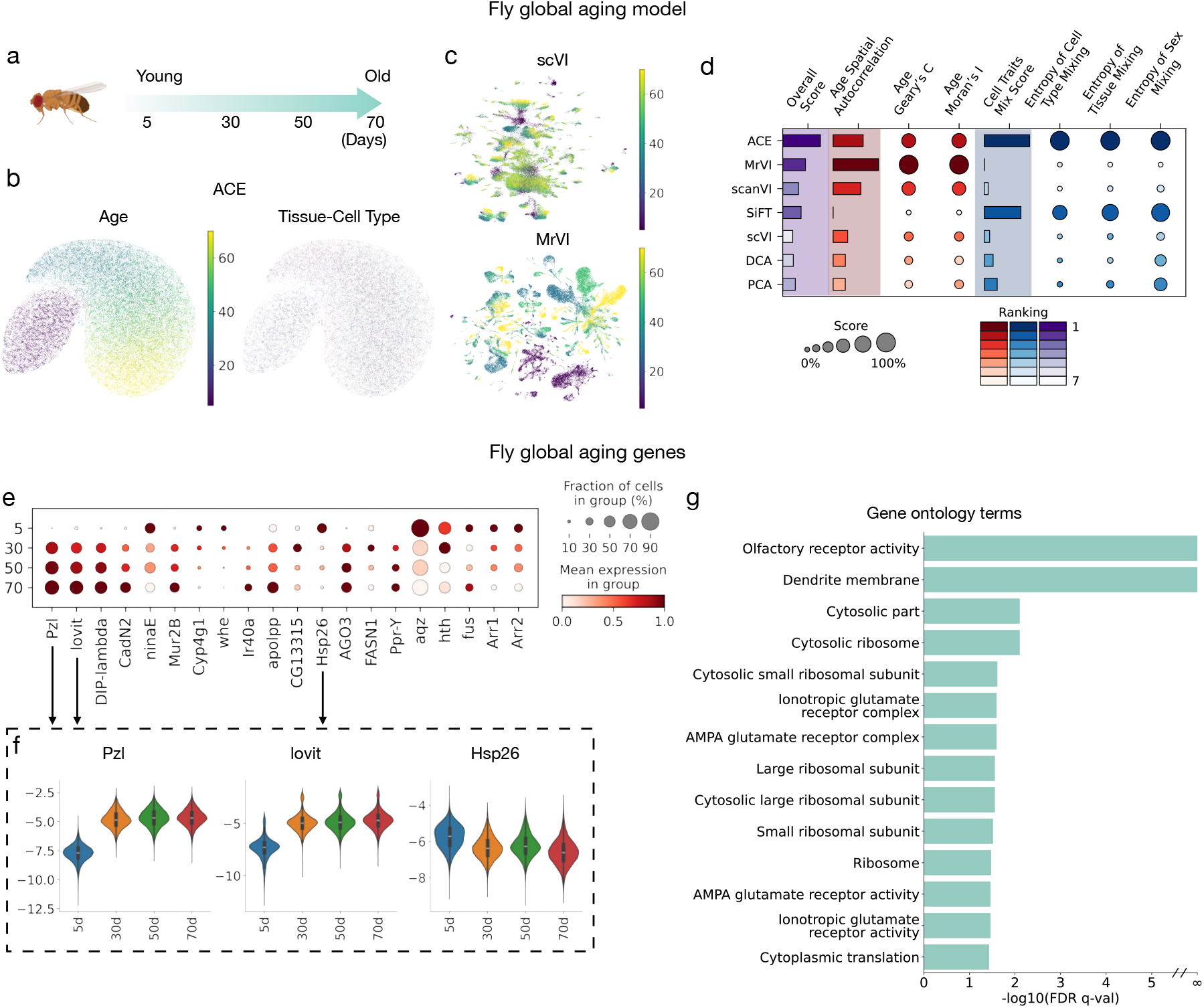
ACE enables global (*i.e*., shared across tissues and cell types) modeling of aging trajectories in fly. **a.** Visualization of the aging timepoints available in the Aging Fly Cell Atlas (AFCA) dataset, ranging from 5 to 70 days, covering the lifespan of a fly. **b**. UMAPs of the age embeddings learned by ACE, colored by age (left) and tissue-cell-type identity (right). The age variables capture aging-related signatures while being disentangled from background factors. **c**. UMAPs of latent variables learned by alternative methods: scVI (top) and MrVI (bottom), colored by age. **d**. Quantitative comparison of ACE and baseline methods (MrVI, scanVI, SiFT, scVI, DCA, and PCA) across multiple evaluation metrics. Circles represent individual metric scores, with circle size indicating the normalized score (scaled between 0 and 1) and color reflecting the relative ranking. For consistency, all metrics are scaled such that higher values indicate better performance. Bars represent composite scores: Age Spatial Autocorrelation is the average of Geary’s C and Moran’s I; Cell Traits Mix Score is the average of cell type, tissue, and sex mixing metrics; and Overall Score is the average of Age Spatial Autocorrelation and Cell Traits Mix Score. Bar color indicates the method’s relative ranking. **e**. Dot plot showing expression patterns of top 20 global aging genes identified by the ACE global aging model in the AFCA. Dot size indicates the fraction of cells expressing the gene in each age group; color represents mean expression. **f**. Violin plots showing expression patterns of representative global aging genes (*lovit, Pzl*, and *Hsp26*) in AFCA data. Violin plots display gene expression distributions across different age groups, aggregated over all tissues and cell types. These visualizations highlight consistent aging-associated expression changes for the selected genes. **g**. Gene set enrichment analysis using the full ranked list of global aging genes from AFCA data identified significantly enriched Gene Ontology terms that play key roles in aging-related biological processes. Significance was assessed at FDR q < 0.05 using the Benjamini-Hochberg correction.

Altogether, the results demonstrate that ACE effectively disentangles global aging signatures from dominant non-aging background factors. Using the XAI method, ACE not only identifies key global aging genes with conserved expression patterns but also uncovers biologically meaningful pathways and gene sets associated with aging. Consistent results across both mouse and fly datasets highlight ACE’s robustness and its ability to reveal aging pathways that are consistent across species at single-cell resolution.

### ACE effectively captures local aging trajectories

While global aging analysis reveals shared, conserved aging signatures, many aging processes are tissue- and cell-type-specific, reflecting the distinct functions and environments of different biological systems. Local (*i.e*., tissue–cell-type–specific) analysis at single-cell resolution is therefore critical for uncovering these context-dependent aging signatures that global models may overlook. As shown in Figure 4a, ACE can also be configured as a local aging model by excluding tissue and cell type from the background factors and using only sex (if available) in the background network. For training the age embedding, we continue to use only the age variable as the predictor. This setup allows the age embeddings to encode both aging-related variation and tissue-cell-type-specific structure, enabling ACE to learn fine-grained, local aging signatures. We applied this configuration to the TMS Droplet dataset, where the resulting UMAPs of ACE’s local age embeddings reveal distinct, coherent trajectories within each tissue–cell-type cluster (Figure 4b,c; Supplementary Figure 10). These results indicate that ACE effectively models tissue-cell-type-specific aging trajectories within a unified framework, without requiring explicit supervision on tissue or cell type

**Figure 4.**
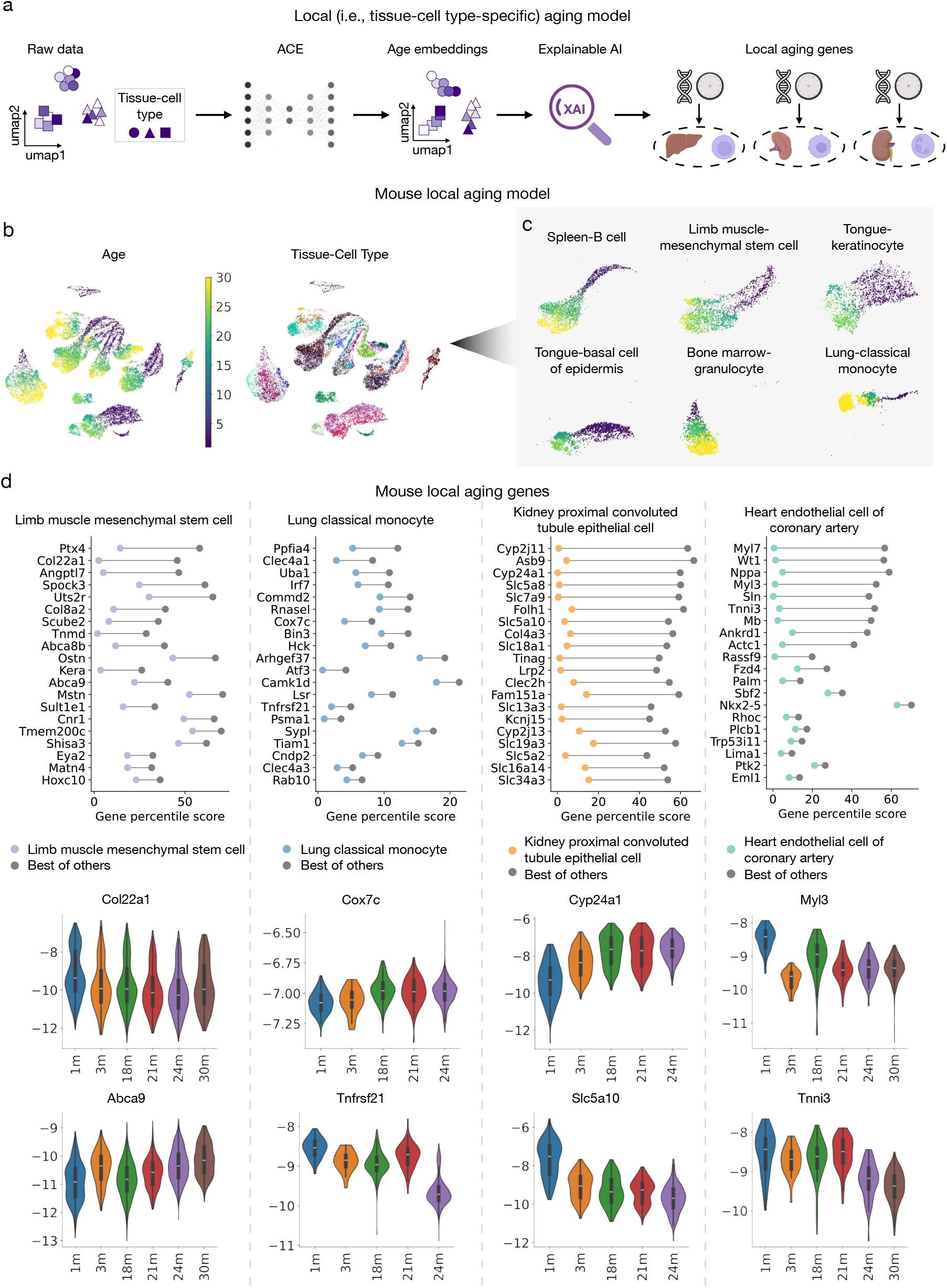
ACE enables local (*i.e*., tissue-cell-type-specific) modeling of aging trajectories. **a.** Overview of the local aging model pipeline. To enable local aging analysis, we train an ACE model while excluding tissue and cell type information from the background factors. This design allows the age embedding to capture both tissue-cell-type-specific clustering and aging-related variation. XAI method, Expected Gradients, is then applied to the ACE model to identify local aging-associated genes specific to tissue-cell-type contexts. **b**. UMAPs of the learned age embeddings, colored by age (left) and tissue-cell-type identity (right). The embedding shows clear clustering by tissue-cell-type while preserving aging trajectories within each cluster. **c**. UMAPs of the age embeddings for examples of tissue-cell-type pairs, illustrating clear aging patterns within individual cell populations. **d**. Local aging genes identified in specific tissue-cell-type pairs in the TMS dataset. **top:** Local aging genes for limb muscle mesenchymal stem cell, lung classical monocyte, kidney proximal convoluted tubule epithelial cell, and heart endothelial cell of coronary artery. The genes are ranked by the difference of their percentile scores within each tissue-cell-type context (colored dots) and their best scores in all other cell types (gray dots). Percentile scores are derived from the full ranked gene list, with lower percentiles corresponding to higher importance. **bottom:** Violin plots showing expression of selected top-ranked local aging genes across age groups, illustrating aging-associated expression trends specific to each tissue-cell-type.

We apply the Expected Gradients (EG) to identify local aging genes for four representative tissue-cell-type pairs, including limb muscle mesenchymal stem cells, lung classical monocytes, kidney proximal convoluted tubule epithelial cells, and heart endothelial cell of coronary artery (Figure 4d, top). For each pair, we calculate the difference between a gene’s percentile score (*i.e*., the relative importance of a gene for a tissue-cell-type pair as estimated by EG) in the target tissue-cell-type and its highest percentile score across all other pairs, and rank the genes by this difference. A high difference indicates that a gene contributes more strongly to aging in one specific tissue-cell-type context while playing a lesser role elsewhere. Many of the top-ranked genes display clear and progressive expression changes across age groups within their respective tissue-cell-type (Figure 4d, bottom). For example, *Col22a1* and *Abca9* in mesenchymal stem cells, *Cox7c* and *Tnfrsf21* in lung classical monocytes, *Cyp24a1* and *Slc5a10* in kidney epithelial cells, and *Myl3* and *Tnni3* in heart endothelial cells of coronary artery, show consistent aging-associated expression patterns. These trends suggest potential roles for these genes in mediating tissue-cell-type-specific aging processes.

Specifically, several of the above-identified genes have been implicated as modulators of tissue-specific pathogenesis. *Col22a1*, a collagen type XXII component expressed in the myotendinous junction, has been identified as a candidate gene for human myopathies [57]. *Abca9* encodes a cholesterol transporter where changes in cholesterol metabolism have been implicated in aging for several tissues [58]. *Cox7c* is a component of the mitochondrial electron transport chain complex IV, where mitochondrial dysfunction is well known to be perturbed in aging [59]. *Tnfrsf21* is a tumor necrosis factor (TNF) receptor superfamily member contributing to inflammatory responses. TNF receptors have been shown to change expression in humans across various ages, indicating aging-dependent remodeling of immune function throughout life [60]. *Cyp24a1* is a major component of vitamin D homeostasis, catabolizing the active form of vitamin D [1,25(OH)2D] into the inactive 1,24,25(OH)2D molecule. Mutations in this gene has been shown to lead to dysregulated vitamin D levels leading to a number of pathological conditions [61, 62]. *Slc5a10* is a kidney-specific SLC transporter that is involved in various carbohydrate reabsorption and has been indicated in diabetic kidney disease [63]. *Myl3* encodes the Myosin light chain 3 protein that has been shown to negatively regulate osteoarthritis in a mouse model, while genetic mutations in this gene have been associated with hypertrophic cardiomyopathy in humans [64, 65]. *Tnni3* encodes troponin I, a component of the myocardial sarcomere structure, where mutations in this structure have been shown to play a role in several cardiomyopathies [66]. The implication of the identified genes in tissue-specific pathologies highlights their potential roles in modulating tissue function across various age stages in humans, suggesting that these genes may be potential targets for therapeutic interventions. We also compared ACE with a previously published linear model Zhang et al. [13] to assess overlap and differences in global and tissue-cell-type-specific aging genes (Supplementary Figure 11).

Overall, ACE not only captures global aging signatures shared across tissues and cell types, but also enables fine-grained analysis of local aging signatures unique to individual tissue-cell-type pairs. This dual capability is essential for uncovering aging mechanisms that are specific to particular biological environments and may be overlooked in global analyses, offering a more comprehensive understanding of the aging process at single-cell resolution.

### ACE enables accurate biological age clocks at both cell and subject levels

ACE’s ability to disentangle aging-related variation from background biological factors enables it to isolate aging-related transcriptomic signatures that reflect the fundamental biology of aging rather than tissue-or cell-type-specific effects. By capturing smooth and continuous aging trajectories, ACE provides a powerful foundation for constructing biological age clocks.

We first assess cell-level biological age prediction using the TMS Droplet mouse dataset. We train a multilayer perceptron (MLP) model that takes ACE-derived age embeddings as input to predict the chronological age of the subject from which each cell was collected (Figure 5a). The model is trained on a randomly selected subset of cells and tested on the remaining held-out cells. The predicted age is defined as the cell’s biological age, reflecting the transcriptomic state of aging. Predicting chronological age from molecular measurements is a widely used approach for estimating biological age [69, 70, 71]. The biological ages closely match the chronological ages (Pearson’s r ≈ 0.9; Figure 5b), with ACE achieving slightly higher correlations than all baseline methods (Supplementary Figure 12a). When evaluated across individual tissue-cell-type pairs, this model achieves strong and statistically significant positive Pearson correlations in nearly all pairs (Figure 5c), with over half of the pairs exceeding a correlation of 0.8. These results demonstrate that ACE effectively captures continuous and generalizable aging signatures, enabling accurate biological age prediction at single-cell resolution.

**Figure 5.**
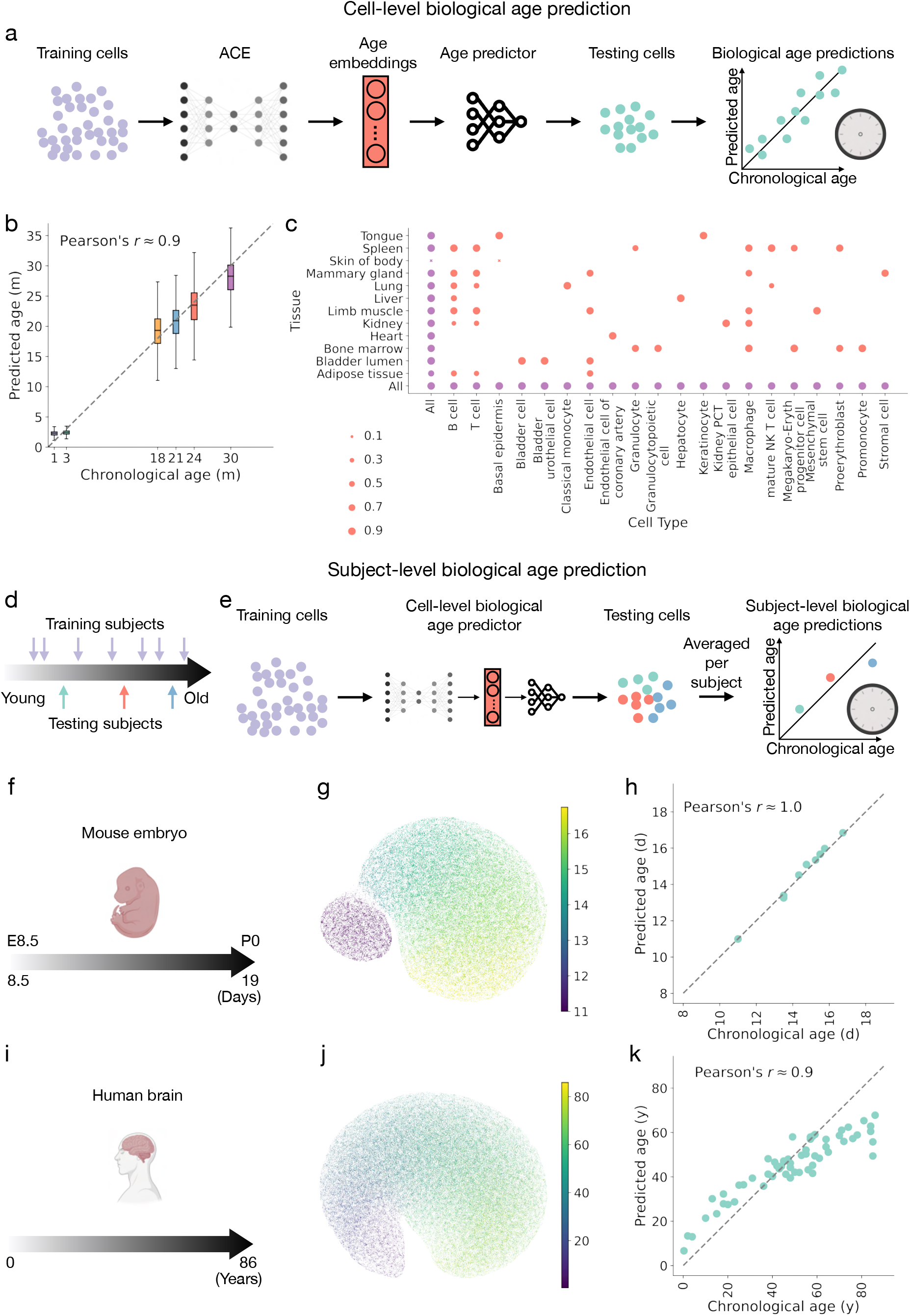
Biological age prediction using age embeddings learned by ACE. **(a-c)** Cell-level aging clock based on ACE embeddings. **a.** Overview of the cell-level aging clock model. ACE learns aging-related latent embeddings from training cells, which are used as input to a multilayer perceptron (MLP) layer for single-cell level age prediction. **b**. Box plots comparing predicted and chronological age across timepoints in TMS Droplet dataset, showing that ACE enables accurate and progressive biological age estimation at the single-cell level. **c**. Dot plot of Pearson correlations between predicted and chronological age across tissue-cell-type pairs. Each dot represents the correlation coefficient for a specific pair, with dot size indicating the strength of the correlation. Cross marks (×) denote negative correlations. All displayed correlations are statistically significant (*p* < 0.05, two-tailed t-test), confirming that the model reliably captures aging-related variation across a broad range of cell type and tissue contexts. **(d-k)** Subject-level aging clock based on ACE embeddings. **d**. Illustration of the subject-level biological age prediction setup. A subset of subjects spanning different ages is held out for testing, while the remaining subjects are used for training. **e**. Overview of the subject-level aging clock model. ACE is trained on cells from the training subjects to learn age embeddings. These embeddings are used to train a cell-level age predictor, which is then applied to cells from the held-out subjects. Predicted cell-level ages are averaged per subject to estimate subject-level biological age. **(f-h)** Developmental age prediction using ACE on a mouse embryo scRNA-seq dataset [67]. The dataset includes 74 subjects sampled between embryonic day 8.5 (E8.5) and postnatal day 0 (P0). Eight developmental time points (11.0, 13.5, 14.333, 14.75, 15.25, 15.5, 15.75, and 16.75 days), each represented by a single subject, are held out entirely from training. **f**. Developmental timeline in the mouse embryo dataset. **g**. UMAP visualization of ACE-derived age embeddings colored by developmental day, illustrating that ACE effectively captures continuous developmental progression. **h**. Predicted versus chronological age for the held-out subjects, demonstrating accurate generalization to unseen developmental stages. **(i-k)** Human brain biological age prediction using ACE. The dataset consists of scRNA-seq profiles from the prefrontal cortex of 286 individuals spanning 0.3 to 86 years of age [68]. A subset of 57 individuals is held out for testing. **i**. Lifespan timeline of the dataset. **j**. UMAP visualization of ACE-derived age embeddings, colored by age at death, illustrating ACE’s ability to learn a continuous aging representation across the human lifespan. **k**. Scatter plot showing predicted versus chronological age at the subject level, indicating strong predictive accuracy across unseen individuals.

We next extend this framework to the subject-level biological age clock, enabling estimation of an individual’s biological age from single-cell profiles. In this setup, we hold out entire test set subjects for evaluation while training ACE and the downstream age predictor on cells from the remaining individuals (Figure 5d,e). For each held-out subject, we apply the trained model to cells from that subject and average the predicted cell-level biological ages to obtain a subject-level biological age estimate. We validate this approach using two distinct datasets: a mouse embryonic development dataset [67] and a human brain aging dataset [68].

The mouse embryo scRNA-seq dataset profiles the transcriptional states of embryos precisely staged at 2-to 6-hour intervals, spanning late gastrulation to birth [67]. After preprocessing, the data we use contains 416,315 cells from 74 embryos spanning embryonic day 8.5 (E8.5) to postnatal day 0 (P0) (Figure 5f; Methods). We train the ACE model using cell type and sex as background factors. Eight randomly selected developmental time points (11.0, 13.5, 14.333, 14.75, 15.25, 15.5, 15.75, and 16.75 days), each corresponding to a single embryo, are completely excluded from training and reserved for testing. ACE captures a smooth developmental trajectory across the held-out time points, generating age embeddings that reflect the known progression of developmental stages (Figure 5g). The predicted ages for the held-out embryos show near-perfect correlation with their true developmental days (Pearson’s r ≈ 1.0; Figure 5h). ACE achieves performance comparable to other deep learning models and clearly exceeds simpler baselines (Supplementary Figure 12b), demonstrating ACE’s ability to generalize across unobserved developmental stages and highlighting the robustness of the ACE-based biological age clock on unseen subjects.

The human brain aging dataset includes scRNA-seq profiles from the dorsolateral prefrontal cortex of postmortem samples spanning the full human lifespan [68]. After preprocessing, the data we use comprises 1,303,449 cells from 286 individuals aged 0.3 to 86 years (Figure 5i; Methods). ACE is trained using cell type and sex as background factors. A randomly selected subset of 57 individuals is entirely held out during training and used for testing. After training, ACE captures a smooth aging trajectory across the human lifespan in the held-out individuals (Figure 5j). Predicted subject-level biological ages show a strong correlation with chronological age (Pearson’s r ≈ 0.9; Figure 5k), with ACE outperforming baseline models (Supplementary Figure 12c), demonstrating its effectiveness for lifespan-wide age prediction from single-cell data and highlighting the generalizability of the ACE-based biological age clock to unseen individuals.

Overall, these findings demonstrate the broad applicability of ACE for biological age prediction at both the single-cell and subject levels. By learning continuous and generalizable age embeddings, ACE supports accurate age estimation across diverse biological scales from individual cells to whole organisms, and performs robustly across various biological contexts. This versatility underscores its potential as a general-purpose framework for biological age estimation across species, tissues, and life stages.

### ACE enables multi-species aging analysis and identification of aging genes conserved across species

Understanding aging across species is crucial for identifying evolutionarily conserved mechanisms that underlie fundamental aspects of the aging process. While many studies focus on organism-specific patterns, learning aging signatures that are shared across species provides an opportunity for discovering *universal* biomarkers and interventions. However, such analyses are challenging due to biological differences and variability in dataset characteristics across species. ACE is well-suited for this task, as its disentangled representation framework enables it to isolate aging signatures from background factors, such as species identity, in multi-species contexts. This allows ACE to extract consistent aging signatures across diverse organisms, positioning it as a unique tool for the analysis of cross-species aging analysis.

To assess whether ACE can uncover conserved aging signatures across species, we extend our analysis to a multi-species setting by integrating three large-scale scRNA-seq datasets: the Tabula Sapiens (TS) human cell atlas [23], the TMS mouse aging cell atlas [9], and the Aging Fly Cell Atlas [10], spanning diverse tissues and life stages (Figure 6a). To enable biologically meaningful comparisons, we map mouse and fly ages to human-equivalent ages using phase equivalencies from established mouse-human mappings [72] and lifespan curve alignment for fly-human mapping (Figure 6b). For humans and mice, we retain the 30 common and most abundant cell types; for flies, we include the 10 most abundant cell types. After preprocessing, the integrated dataset comprises 520,139 human cells, 171,924 mouse cells, and 101,684 fly cells across 19 tissues and 40 cell types (see Supplementary Figure 13 for the UMAPs of the raw data; Methods). The inclusion of mouse and fly data substantially broadens the age distribution compared to human data alone (Figure 6b). ACE is then trained on the multi-species dataset using 2,515 one-to-one orthologous genes shared across all species, with species, tissue, cell type, and sex specified as background factors.

**Figure 6.**
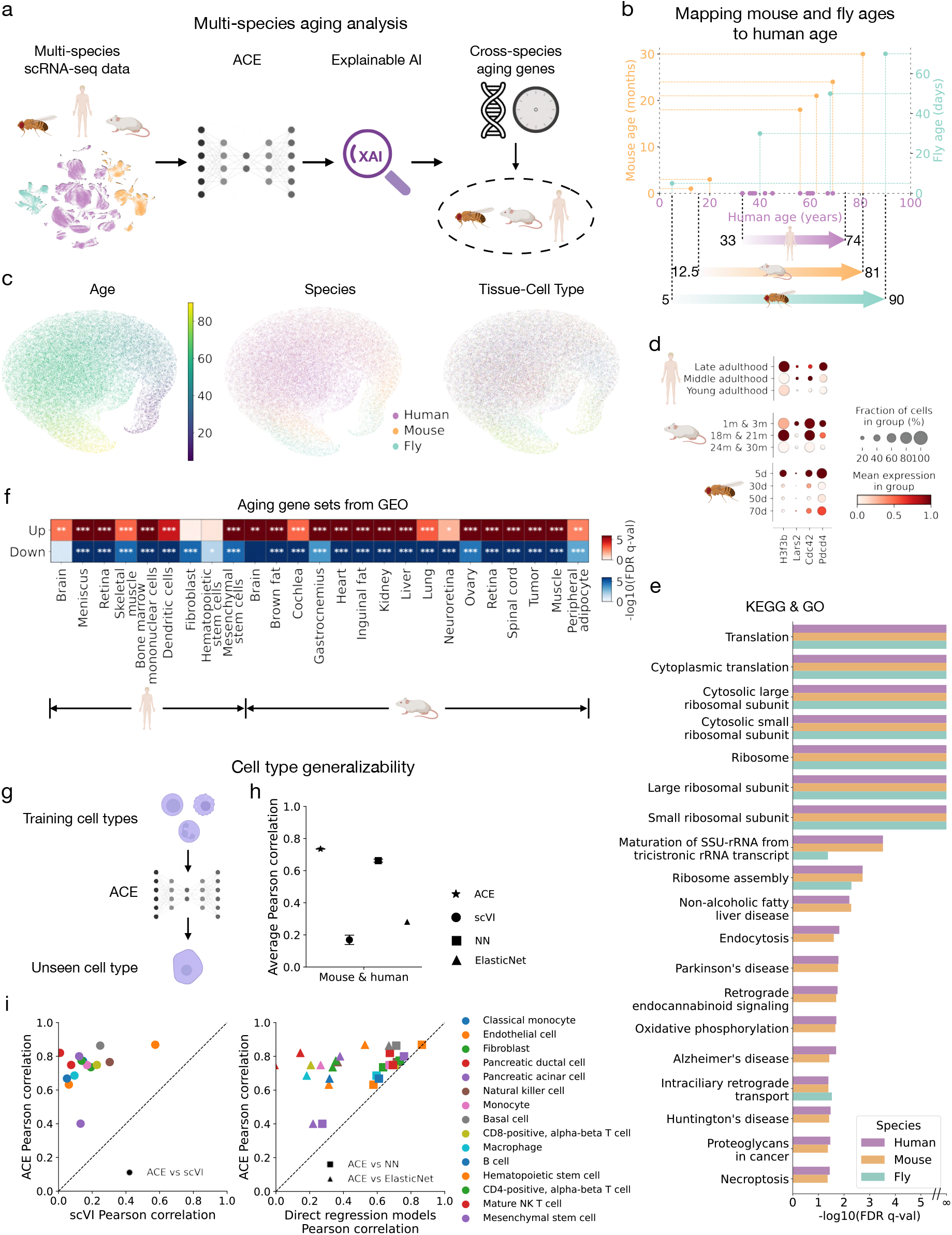
Multi-species aging analysis and cell type generalizability using ACE. **a.** Overview of the multi-species aging analysis pipeline. ACE is applied to multi-species scRNA-seq data from human, mouse, and fly to disentangle aging-related signatures shared across species. XAI method, Expected Gradients, is then used to identify conserved aging genes shared across species. **b**. Mapping of mouse and fly ages to human-equivalent ages using lifespan alignment, enabling biologically meaningful cross-species comparisons. **c**. UMAPs of the age embeddings learned by ACE from integrated human, mouse, and fly data. Embeddings are colored by age (left), species (middle), and tissue-cell-type labels (right), showing that the model captures a smooth aging trajectory across species, with cells from different organisms well mixed in the age latent space. **d**. Expression patterns of selected cross-species aging-associated genes identified by ACE. Shown are normalized expression levels of orthologous genes across different age groups in human, mouse, and fly, labeled using mouse gene symbols (*H3f3b, Lars2, Cdc42*, and *Pdcd4*). Dot size indicates the fraction of cells expressing the gene; color represents mean expression. **e**. KEGG and GO pathway enrichment analysis of conserved aging genes across human, mouse, and fly identified conserved pathways that are significantly enriched and involved in key aging-related biological processes. Significance was assessed at FDR *q* < 0.05 using the Benjamini-Hochberg correction. **f**. Gene set enrichment analysis using the “Aging Perturbations from GEO Up” and “Aging Perturbations from GEO Down” gene set collections. These databases consist of gene sets curated from studies in GEO that compare aged versus young samples, capturing genes consistently upregulated or downregulated with age for various tissues and cell types. The results demonstrate that multi-species ACE-derived aging gene rankings are significantly enriched in multiple known human and mouse aging-associated gene sets across various tissues and cell types. Red boxes indicate enrichment in *upregulated* gene sets; blue boxes indicate enrichment in *downregulated* gene sets. Color intensity reflects — log_10_(FDR *q*-value), with significance determined by FDR-adjusted *q*-values using the Benjamini-Hochberg correction. Asterisks denote significance thresholds (^∗^ *p* < 0.05, ^∗∗^ *p* < 0.01, ^∗∗∗^ *p* < 0.001). **g**. Schematic illustrating the evaluation of cell type generalizability. To assess how well age embeddings learnt by ACE generalize to new biological contexts, the model is trained on mouse and human data excluding one cell type. Age embeddings are then extracted for the held-out (unseen) cell type, and used to train a multilayer perceptron (MLP) for age prediction. This setup evaluates how effectively the learned age embeddings capture aging signatures that generalize across cell types. **h**. Average Pearson correlation between predicted and true age across unseen cell types in mouse and human, comparing ACE to baseline models including scVI, a neural network (NN), and ElasticNet. Age prediction is performed using different inputs and models: MLPs are trained on age embeddings from ACE, latent variables from scVI, and raw gene expression for NN; ElasticNet is trained directly on raw gene expression. Each dot represents the mean Pearson correlation across 10 runs with different random seeds (for ACE, scVI, and NN); error bars indicate the standard deviation. ACE consistently outperforms baseline models. **i**. Scatter plots comparing ACE to baseline models across individual cell types, demonstrating that ACE provides stronger generalization to unseen cell types.

The learned age embeddings reveal a smooth and continuous aging trajectory shared across all species, with cells from different organisms, tissues, and cell types well mixed in the same embedding space (Figure 6c; see Supplementary Figure 14 for UMAPs of both age and background embeddings colored by age and background factors). We then apply the Expected Gradients to identify aging-associated genes that are consistently important across species. Several conserved genes, such as *H3f3b, Lars2, Cdc42*, and *Pdcd4*, exhibit progressive and consistent expression changes across age groups in all species (Figure 6d). The conserved genes across species all play a role in known aging-related processes. *H3f3b* encodes a histone protein involved in chromatin structure and epigenetic regulation of gene expression, which is a known pathway involved in aging [6]. *Lars2* encodes a mitochondrial leucyl-tRNA synthetase protein. A study utilizing *C. elegans* demonstrated that downregulation of this gene was associated with longer lifespan [73]. *Cdc42* encodes a GTPase of the Rho-superfamily that regulates the actin cytoskeleton, cell cycle, and proliferation of many cell types and has been implicated in numerous studies as a factor in aging [74]. *Pdcd4* is involved in regulating programmed cell death and the knockdown of this gene in hepatoma cancer cells has been shown to induce cellular senescence [75].

Pathway enrichment analysis using KEGG and GO shows that these conserved aging genes are significantly enriched for translation-related functions (*e.g*., ribosome, cytoplasmic translation) across all 3 species, highlighting the importance of proteostasis as a conserved aging mechanism across species. A greater number of conserved pathways are present between the human and mouse datasets, including immune signaling (*e.g*., endocytosis), metabolic processes (*e.g*., oxidative phosphorylation), and neurodegenerative pathways (*e.g*., Alzheimer’s and Parkinson’s diseases) (Figure 6e; see Supplementary Figure for the full list of enriched pathways). Furthermore, ACE’s ranked gene list demonstrates strong and consistent enrichment across nearly all aging-associated gene sets curated from GEO for both human and mouse, spanning multiple tissues and cell types (Figure 6f). These results support its ability to identify aging-related genes that are shared across species, tissues, and cell types.

Because ACE captures aging signatures shared across diverse biological contexts, it should also generalize to new, unseen contexts. To evaluate this, we test ACE’s ability to generalize to previously unseen cell types. Using a combined dataset of mouse and human cells across 40 common cell types, we select 15 with sufficient cell numbers and age diversity to serve as held-out cell types. For each held-out evaluation, we train ACE on the 39 cell types and assess its age prediction performance on the excluded one (Figure 6g). An MLP is trained on the ACE-derived age embeddings to predict age. We compare ACE to baseline methods including scVI, which uses its learned embeddings with an MLP for age prediction, as well as a neural network (NN) and an ElasticNet model trained directly on raw gene expression for age prediction. Across the 15 held-out cell types, ACE achieves an average Pearson correlation of 0.73, with 11 cell types exceeding 0.7 (Figure 6h), representing a 10.6% improvement over the best alternative baseline, NN. ACE consistently outperforms all baselines, demonstrating its ability to generalize aging predictions to previously unseen cell types and supporting its utility in modeling shared aging signatures across heterogeneous biological contexts (Figure 6h,i).

Together, these findings demonstrate ACE’s ability to model aging trajectories across diverse organisms and uncover conserved aging signatures, while also generalizing to previously unseen cell types. By learning shared aging representations that disentangle background biological factors, ACE enables robust aging analysis across species, tissues, and cell types. This highlights its versatility as a unified framework for cross-species and cross-context aging studies at the single-cell resolution.

### Experimental Validation of ACE-Identified Aging Genes

To assess the biological relevance of aging-related genes identified by ACE, we performed RNAi knockdown experiments in *C. elegans* and measured their effects on lifespan (Figure 7; Methods). We focused on genes identified by the mouse global aging model, as the mouse aging dataset provides a comprehensive, aging-focused single-cell resource that yields less confounded aging signatures. Specifically, we selected 8 mouse genes from the top 150 most important global aging genes identified by the mouse global aging model: *Ppia, Ftl1, Gpx3, Rpl37, Serpina3k, Tpt1, Rps27a*, and *Uba52*. Mouse genes were mapped to their *C. elegans* orthologs, resulting in a total of 15 tested *C. elegans* genes. These genes were selected based on their potential involvement in proteostasis, their evolutionary conservation across species, and the feasibility of functional testing in *C. elegans* via RNAi, as they do not produce larval arrest or lethality phenotypes.

**Figure 7.**
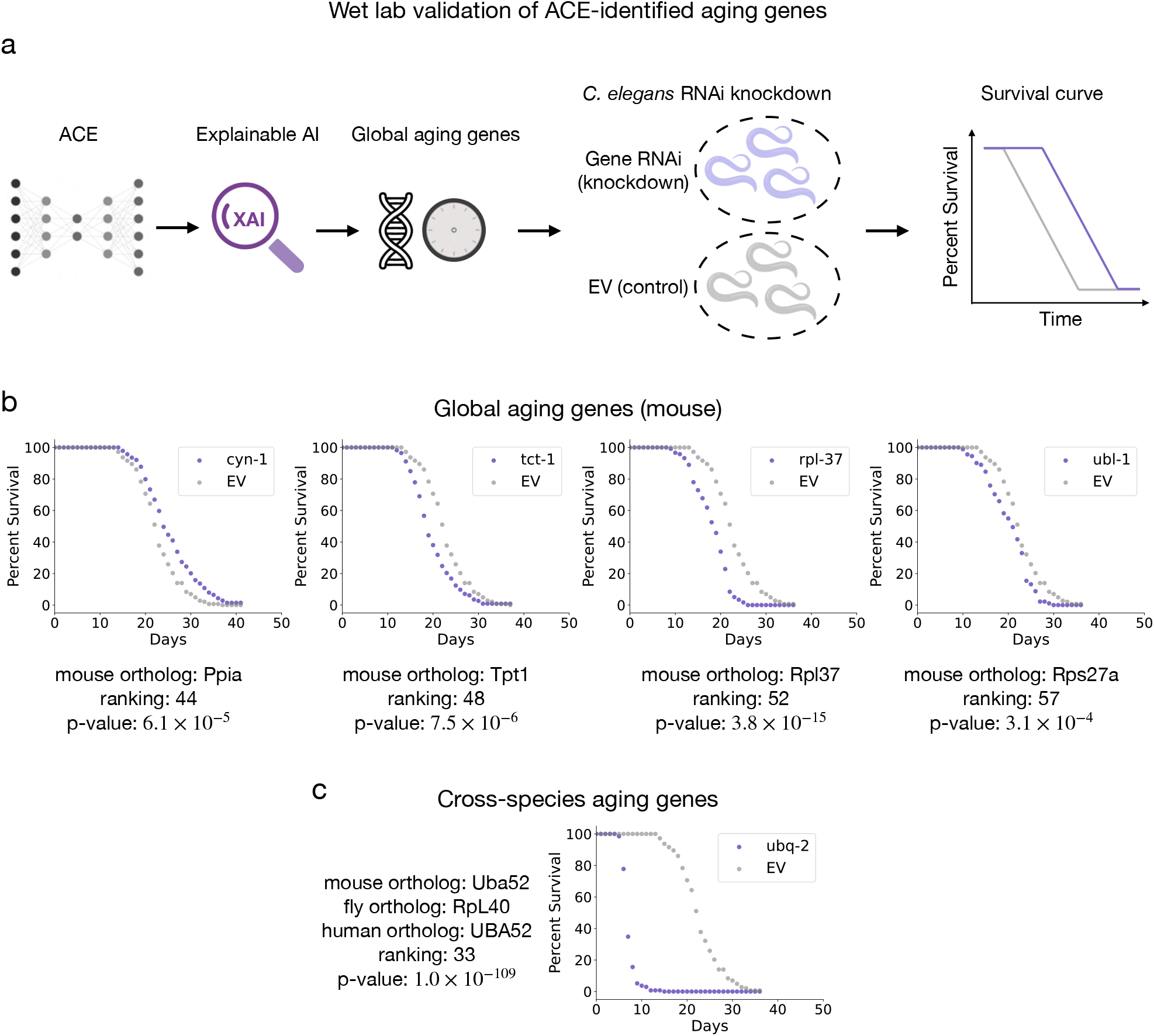
Wet lab validation of ACE-identified aging genes using C. elegans RNAi lifespan assays. **a.** Overview of the experimental validation pipeline. ACE identifies global aging genes using XAI techniques. Selected genes are knocked down in *C. elegans* via RNA interference (RNAi), and survival is compared to empty vector (EV) controls. **b**. Survival curves for knockdown of *C. elegans* orthologs corresponding to selected global aging genes identified by the mouse global ACE model. Each plot shows the percent survival of animals treated with gene-specific RNAi versus empty vector control. Mouse gene names, gene rankings from the mouse global ACE model, and p-values from t-tests (adjusted using the Benjamini-Hochberg correction) are shown below each plot. **c**. Survival curves for knockdown of *C. elegans* genes corresponding to a top-ranked aging-associated gene from the cross-species ACE model. Orthologs are selected based on mouse gene mappings. The plot shows percent survival for animals treated with gene-specific RNAi versus EV. Mouse gene name, its one-to-one orthologs in human and fly, gene rankings from the cross-species ACE model, and adjusted p-value from t-tests (adjusted using the Benjamini-Hochberg correction) are shown below each plot.

For each gene, *C. elegans* orthologs were knocked down using RNAi, and survival curves were compared to those of empty vector (EV) controls. As shown in Figure 7b, four knockdowns, *cyn-1, tct-1, rpl-37*, and *ubl-1*, led to significant lifespan alterations, with p-values ranging from 3.1 × 10^−4^ to 3.8 × 10^−15^, confirming that ACE effectively prioritizes functionally relevant aging genes. Importantly, three of these four genes (*cyn-1, rpl-37, ubl-1*) are involved in protein translation. These results corroborate the cross-species pathway analysis (Figure 6e), further emphasizing the importance of proteostasis as a conserved aging mechanism across species. Of the 8 mouse genes tested, 6 had at least one *C. elegans* ortholog whose knockdown significantly impacted lifespan. Among the 15 *C. elegans* genes tested, 8 showed significant effects (see Supplementary Figure 18 for full survival curves).

We also observed that one of the tested genes, *Uba52*, ranked highly in the cross-species ACE model (ranked 33), suggesting its conserved relevance across mouse, human, and fly. As shown in Figure 7c, knock-down of *ubq-2* (the *C. elegans* ortholog of *Uba52*) led to significantly reduced lifespan, further supporting the biological validity of ACE’s prioritizations across species. These results highlight ACE’s ability to uncover evolutionarily conserved regulators of aging that can be experimentally validated in model organisms.

These experimental validations demonstrate ACE’s power in prioritizing biologically meaningful aging-associated genes, with a strong emphasis on proteostasis. The observed lifespan effects in *C. elegans* following knockdown of ACE identified genes highlight the model’s functional relevance and cross-species generalizability. Notably, several of the validated genes play key roles in ribosomal function and ubiquitin-mediated protein turnover – hallmarks of proteostasis that are increasingly recognized as central to aging. These findings underscore ACE’s potential to uncover evolutionarily conserved regulators of aging and provide mechanistic insights with broad implications for aging biology and translational geroscience.

## Discussion

ACE provides an interpretable and generalizable framework for disentangling aging-related signatures from background biological variation in single-cell transcriptomic data. By modeling aging trajectories across tissues, cell types, and species, ACE enables the identification of both global and local aging signatures, revealing how aging manifests in conserved and context-dependent ways. Applied to large-scale mouse, fly, and human atlases, ACE uncovered robust, biologically meaningful aging-associated genes and pathways, including well-studied aging genes such as *Sparc* [76], *S100a8* [77], and *Hsp26* [78, 79]. ACE’s learned embeddings also enabled accurate subject-level biological age prediction in mouse embryo and human brain datasets, highlighting their utility for biological age estimation.

Across species, our results overwhelmingly implicated protein homeostasis in aging. Protein homeostasis, or proteostasis, incorporates several processes, including protein synthesis, maintenance, and destruction/turnover in cells and has long been implicated in the aging process [80]. ACE has identified critical conserved components of proteostasis that affect aging across several species, including several genes involved in ribosome and ubiquitination, which are cellular machinery involved in protein synthesis and protein degradation, respectively. Importantly, several of these ribosomal and protein ubiquitination genes identified in the mouse model and multi-species model were validated in a *C. elegans* model and were found to significantly impact lifespan. The validation of these genes in an animal model unique from the animal models used in the ACE training data highlights the importance of these processes as conserved aging mechanisms across species. Additionally, this highlights the generalizability of ACE towards other model organisms and its broad biological utility. The identification of proteostasis genes is consistent with other experimental results showing that long-lived clams and long-lived naked mole rats have exceptional proteostasis compared to other similar species (shorter-lived clams and rats), further implicating proteostasis as foundational in the aging process. Indeed, it seems logical that cells need to maintain the function of the proteins that produce, repair and maintain all aspects of cellular physiology. Together, these results support the utility of ACE to ascertain genes and pathways involved in aging from publicly available scRNA-seq datasets.

The potential for ACE to identify previously unidentified driver genes involved in cellular aging is an important use-case for the model. With an increasingly aging population, the identification of aging driver genes can allow for the identification of druggable targets that can help ease the burden of aging and improve healthy lifespan. The application of ACE to identify tissue- and cell type-specific aging genes can assist with the therapeutic intervention for tissue-specific aging-associated diseases, such as aging-associated neurodegeneration, cardiovascular disease, and chronic inflammatory diseases. Genes identified using ACE can be studied in commonly used animal models, as performed here, or they can be studied in human-specific models such as primary cell lines or human induced pluripotent stem cell (hiPSC) derived cell lines and organoids. The utilization of an AI model like ACE in combination with human-derived cell models is in direct line with the 2022 FDA Modernization Act 2.0, which allows for the use of new approach methodologies (NAMs) as an alternative to animal models for use in safety and efficacy assessments in the drug development process [81]. Models like ACE could prove to be a powerful tool in the drug discovery phase during preclinical drug development.

From a technical perspective, ACE introduces an interpretable and modular generative framework that disentangles aging-related variation from complex background factors in scRNA-seq data. Unlike conventional models that compress all variation into a single latent space [15, 29, 30], ACE learns two embeddings to isolate aging signatures, improving both interpretability and generalization. Its flexible design extends naturally to other biological settings, enabling disentanglement of molecular signatures linked to diverse phenotypes such as neurodegeneration, tumor progression, and immune activation. ACE also lays the groundwork for developing single-cell aging foundation models that generalize across cell types, tissues, species, and experimental conditions. Unlike existing foundation models trained on heterogeneous datasets that may conflate aging with other biological effects [82, 83, 84], ACE explicitly disentangles aging signa-tures from background variation, offering a targeted and interpretable method for building aging-specific foundation models. This disentanglement principle further positions ACE as a framework for developing phenotype-specific foundation models in single-cell biology.

The limitation of ACE is that it currently relies on accurate annotations for background factors such as tissue and cell type, which may be incomplete or inconsistent across datasets. This dependence could affect model performance on less curated or cross-study data. In addition, ACE currently operates on transcriptomic data only. Incorporating additional omics layers, such as DNA methylation [85], chromatin accessibility [86], or proteomics [87], could provide a more comprehensive understanding of aging. Advances in multi-modal generative modeling [17, 88] present promising opportunities to extend ACE toward multiomics integration, thereby enhancing its biological scope.

In summary, ACE enables interpretable and disentangled modeling of aging signatures at single-cell resolution across species, tissues, and cell types. Its scalability and generalizability make ACE a powerful framework for uncovering biologically meaningful signatures from high-dimensional single-cell data and for advancing phenotype-specific modeling in aging and disease research.

## Methods

### The ACE model

Here, we present the ACE model in more detail. We begin by describing the model’s generative process and then the model’s inference procedure.

#### The ACE generative process

For a data point *x*_*n*_, we assume that each expression value *x*_*ng*_ for sample n and gene g is generated through the following process:

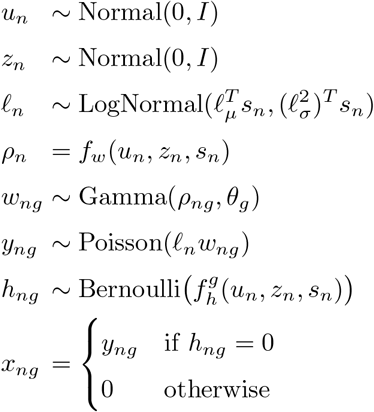

Here, *u*_*n*_ and *z*_*n*_ denote two sets of latent variables that account for variations in scRNA-seq expression data. Specifically, *u*_*n*_ is designed to capture variation primarily driven by age (*a*_*n*_), aiming to isolate aging-related biological signatures. In contrast, *z*_*n*_ models background variation associated with covariates *b*_*n*_ (e.g., cell type, tissue, sex, species), which are often dominant sources of variation and may overshadow subtle aging signatures. ACE’s design enables the disentanglement of these two factors, helping to recover aging-related signatures across complex biological contexts. We place standard multivariate Gaussian priors on both *u*_*n*_ and *z*_*n*_ to support efficient inference within the variational autoencoder (VAE) framework [89].

To ensure that the latent variables *u*_*n*_ and *z*_*n*_ accurately capture distinct sources of biological variation, specifically aging-related signatures and background-related signatures, we adopt two complementary strategies. First, during training, we add supervised prediction losses that encourage each latent variable to retain information about its corresponding covariates. Specifically, a age prediction network is trained to predict the chronological age of the donors (*a*_*n*_) from *u*_*n*_, while a background prediction network is trained to predict background covariates *b*_*n*_ (e.g., cell type, tissue, sex, species) from *z*_*n*_. These covariates can be binary, categorical, or continuous; accordingly, we apply cross-entropy loss for classification tasks and mean squared error for regression. Second, we apply the d-variable Hilbert-Schmidt Independence Criterion (dHSIC) [25, 90, 26, 24] to enforce statistical independence between *u*_*n*_ and *z*_*n*_. Since the dHSIC equals zero if and only if the variables are independent, minimizing this quantity ensures that the two latent spaces encode non-overlapping biological signatures, thereby disentangling aging-related variation from dominant background effects.

Here ℓ_*µ*_ and 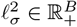, where *B* denotes the cardinality of an optional label denoting experimental batch, parameterize the prior for a latent RNA library size scaling factor on a log scale, and *s*_*n*_ is a *B*-dimensional one-hot vector encoding the batch label for each cell. For each batch, ℓ_*µ*_ and 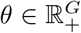 are set to the empirical mean and variance of the log library size. ρ_*ng*_ ∈ ℝ_+_ and shape θ_*g*_ ∈ ℝ_+_ parameterize our Gamma distribution with a mean-shape parameterization. Moreover, we note that θ_*g*_ can be viewed as a gene-specific inverse dispersion parameter for a negative binomial distribution, and we learn 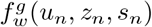 through variational inference. *f*_*w*_ and *f*_*g*_ are neural networks that transform the latent space and batch annotations to the original gene space, i.e. *f*_*w*_: ℝ^*d*^ × *{*0, 1*}*^*B*^ → ℝ^*G*^ and similarly for *f*_*g*_, where d is the combined dimensionality of the concatenated age and background latent spaces. The outputs of the network *f*_*w*_ represent the mean proportion of transcripts expressed across all genes, and this constraint is enforced by using a softmax activation function in the last layer. That is, letting 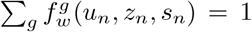 denote the entry in the output of *f*_*w*_ corresponding to gene *g*, we Have 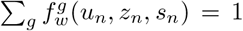. The output of the neural network *f*_*h*_ denotes whether a dropout event has occurred (i.e., a gene’s expression is read as zero due to technical factors rather than meaningful biological phenomena).

Our generative process closely follows that of contrastiveVI [21], but with the notable additions of prediction networks and the dHSIC penalty. While contrastiveVI’s modeling approach excels situations when explicit case and control groups of cells are available, it is not appropriate for scenarios without a clear control group. By incorporating age covariate a_*n*_ and background covariates *b*_*n*_, and introducing prediction networks for both, ACE can effectively isolate the aging variations even without an explicit group of corresponding control cells.

### Inference with ACE

We cannot compute the ACE posterior distribution using Bayes’ rule as the integrals required to compute the model evidence *p*(*x*_*n*_|*s*_*n*_) are analytically intractable. As such, we instead approximate our posterior distribution using variational inference [91]. We approximate our posterior with a distribution factorized as follows:

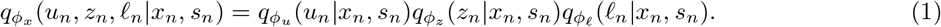

Here *ϕ*_*x*_ denotes a set of learned weights used to infer the parameters of our approximate posterior. Based on our factorization of the posterior in Eq. 1, we can divide our full set of parameters *ϕ*_*x*_ into disjoint subsets *ϕ*_*u*_, *ϕ*_*z*_ and *ϕ*_*A*_ for inferring the parameters of the distributions of *u, z* and ℓ respectively. As in the VAE framework [89], we approximate the posterior for each set of latent variables via a deep neural network that takes in expression levels as input and returns the parameters of its corresponding approximate posterior distribution. Moreover, we note that each factor in the posterior approximation shares the same family as its respective prior distribution (e.g. *q*(*u*_*n*_|*x*_*n*_,*s*_*n*_) follows a normal distribution). By marginalizing out *<u>w</u>*_*ng*_, *h*_*ng*_, and *y*_*ng*_, we can simplify our likelihood yielding *p*_*ν*_(*x*_*ng*_|*u*_*n*_, *z*_*n*_, *s*_*n*_, *ℓ*_*n*_), which has a closed form of a zero-inflated negative binomial (ZINB) distribution and where ν denotes the parameters of our generative model. We implement our generative model with deep neural networks as done for our approximate posterior distributions. For Eq. 1, We derive the following Evidence Lower Bound (ELBO) for the marginal log-likelihood:

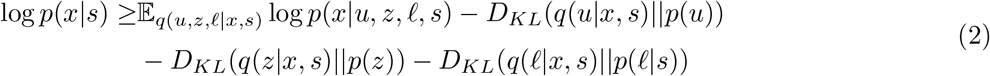

We optimize the parameters of the generative model, inference networks, and prediction networks jointly using stochastic gradient descent. The optimization objective is a weighted composite loss that combines the ELBO, prediction losses for the age covariate *a*_*n*_ and background covariate *b*_*n*_, and a penalty term based on the *d*HSIC, which encourages disentanglement between *u*_*n*_ and *z*_*n*_. The neural networks used for both the variational and generative distributions are standard feedforward architectures with typical activation functions. Formally, the total loss is defined as:

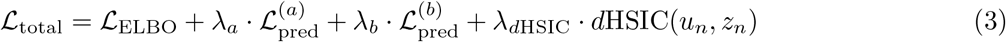

where, ℒ_ELBO_ denotes the negative ELBO as defined in Eq. 2; 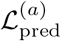 is the prediction loss for the chronological age *<u>a</u>*_*n*_, for which we use mean squared error (MSE) since age is a continuous variable; 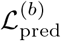 represents the prediction loss for the background covariates *b*_*n*_, such as cell type, tissue, sex, and species, and we apply cross-entropy loss due to their categorical nature; *d*HSIC(*u*_*n*_, *z*_*n*_) quantifies the statistical dependence between the latent variables *u*_*n*_ and *z*_*n*_. The hyperparameters *λ*_*a*_, *λ*_*b*_, *λ*_*d*HSIC_ are used to scale the auxiliary loss components so that their magnitudes are comparable to that of *L*_ELBO_. This normalization facilitates stable joint optimization by preventing any single auxiliary loss from dominating the training dynamics.

### Model optimization details

For all datasets, ACE models were trained using 64% of cells, validated on 16% to determine the optimal number of training epochs, and evaluated on the remaining 20% for both quantitative and qualitative analyses. Early stopping was applied when the validation ELBO failed to improve for 45 consecutive epochs. All models were optimized using the Adam optimizer with a learning rate of 0.0001, ϵ = 0.01, and a weight decay of 10^—6^. The hyperparameters λ_*a*_, λ_*b*_, λ_*d*HSIC_ were chosen separately for each model to ensure that auxiliary loss terms were on a comparable scale to the ELBO.

Each ACE model was trained with 20 latent dimensions: 3 reserved for aging-related variation and 17 for background variation, unless otherwise noted. For the mouse and fly global aging models, and the mouse embryo model, background covariates included cell type, tissue, and sex. The human brain aging model used cell type and sex as background covariates. For the mouse local aging model, background included only sex, and 13 age dimensions and 7 background dimensions were used to encourage learning tissue- and cell type-specific aging signatures. For the multi-species aging model, background covariates included cell type, tissue, sex, and species.

We compared ACE with the following baseline methods: MrVI [29], scANVI [30], SiFT [31], scVI [15], DCA [16], and PCA. Each baseline was trained with 20 latent dimensions for consistency. For MrVI, chronological age was used as the “sample key” to estimate age effects. For scANVI, we treated age as a categorical “cell type label” for conditional modeling. For SiFT, we treated cell type, tissue, and sex as unwanted variation sources and removed them accordingly during training.

### Datasets and preprocessing

We now briefly describe all datasets used in this work along with any corresponding preprocessing steps. All preprocessing steps were performed using the `Scanpy` Python package.

#### Tabula Muris Senis mouse dataset

The Tabula Muris Senis (TMS) dataset [9] is a large-scale single-cell RNA-sequencing (scRNA-seq) atlas characterizing aging-associated transcriptomic changes across multiple tissues in the mouse. It consists of two branches based on sequencing protocols: TMS Droplet, generated using microfluidic droplet-based platforms, and TMS FACS, produced via fluorescence-activated cell sorting (FACS) followed by Smart-seq2 library preparation. In our study, we used the TMS Droplet dataset as the primary resource due to its larger number of cells and used the TMS FACS dataset to validate key biological findings.

#### Preprocessing for TMS Droplet

We first filtered out genes expressed in fewer than 5 cells and removed cells expressing fewer than 500 genes. Cells with fewer than 3,000 total counts were also excluded. To ensure sufficient representation, we retained only the top 20 most abundant cell types and included tissues with more than 1,000 cells. We then selected the top 2,000 highly variable genes using the Seurat v3 method with batch correction applied via the `batch` key, based on raw counts. In addition to these HVGs, we incorporated aging-associated genes identified in the study by Zhang et al. [13].

After preprocessing, the TMS Droplet dataset comprised 135,420 cells and 4,327 genes, derived from 17 male and 6 female mice across six age groups (1m, 3m, 18m, 21m, 24m, 30m), 13 tissues, and 20 cell types.

#### Preprocessing for TMS FACS

We applied similar preprocessing steps to the TMS FACS dataset. Specifically, we removed cells expressing fewer than 500 genes and excluded those having fewer than 3,000 total counts. We then filtered to retain only tissues and cell types that were also present in the TMS Droplet data. Furthermore, we removed cell types, tissues, and donors with fewer than 100 cells, and limited the dataset to age groups with more than 1,000 cells. We restricted the gene set to those retained in the processed TMS Droplet data for consistency.

The final TMS FACS dataset comprised 24,546 cells and 4,120 genes across three age groups (3m, 18m, and 24m), including samples from 12 male and 6 female mice.

#### Aging Fly Cell Atlas

The Aging Fly Cell Atlas [10] is a large-scale single-cell RNA-sequencing resource that profiles aging in *Drosophila melanogaster* across multiple tissues, time points, and sexes. It provides comprehensive coverage of aging-associated transcriptional changes in both neural and peripheral tissues, offering insights into conserved and fly-specific aging processes.

We performed the following preprocessing steps. First, we removed genes expressed in fewer than 5 cells and filtered out cells expressing fewer than 300 genes. Cells with fewer than 500 total counts were also excluded. To ensure sufficient representation, we retained only cell types with more than 1,000 cells and excluded cells with ambiguous sex annotations labeled as “mix.” We then selected the top 2,000 highly variable genes using the Seurat v3 method on the raw count layer with batch correction.

After preprocessing, the final dataset consisted of 424,863 cells and 2,000 genes, covering 14 cell types across two tissues (head and body), four age groups (5d, 30d, 50d, and 70d), and two sexes.

#### Mouse embryo dataset

To evaluate ACE for biological age prediction, we used the mouse embryo single-cell RNA-sequencing dataset [67]. This dataset provides high-resolution temporal profiling of developing mouse embryos, covering time points from embryonic day (E) 8.5 to postnatal day 0 (P0), sampled every 2 to 6 hours. It enables modeling of developmental trajectories as a continuous biological aging process.

We first filtered out genes expressed in fewer than 5 cells and removed cells expressing fewer than 500 genes. Cells with fewer than 3,000 total counts were also excluded. To ensure sufficient representation, we retained only the top 20 most abundant cell types. We then selected the top 2,000 highly variable genes using the Seurat v3 method, based on raw count values.

After preprocessing, the final dataset consisted of 416,315 cells from 74 embryos, spanning the developmental window from E8.5 to P0. For training, we modeled developmental stage as the aging covariate and treated cell type and sex as background covariates. To assess the model’s ability to generalize to unseen time points, we excluded eight developmental stages, E11.0, E13.5, E14.333, E14.75, E15.25, E15.5, E15.75, and E16.75, from training. Each excluded time point corresponds to a single embryo and was reserved for testing.

#### Human brain aging Dataset

To evaluate biological age prediction in the human brain, we utilized the dorsolateral prefrontal cortex (dlPFC) single-cell RNA-sequencing dataset [68]. This dataset provides a comprehensive transcriptomic atlas of the human lifespan in a critical cognitive brain region.

We selected the top 2,000 highly variable genes using the Seurat v3 method, based on raw count values. After preprocessing, the final dataset consisted of 1,303,449 cells from 286 individuals aged between 0.3 and 86 years, spanning eight major cell types. To enable unbiased evaluation, a randomly selected subset of 57 individuals was entirely held out from training and used exclusively for testing.

#### Multi-species dataset (Mouse + Fly + Human)

To enable comparative analysis of aging across species, we constructed a multi-species dataset by combining the TMS mouse data, the Aging Fly Cell Atlas, and the Tabula Sapiens (TS) human cell atlas [23]. Tabula Sapiens is a comprehensive human single-cell transcriptomic atlas spanning multiple organs and systems from healthy adult donors, serving as a reference for human cellular diversity.

To align gene expression features across species, we first identified one-to-one orthologs between mouse and human, and between fly and human, retaining only genes with one-to-one orthology in both comparisons. We then merged the datasets across species based on the shared orthologous gene set.

We performed standard filtering to ensure quality and comparability: genes expressed in fewer than 5 cells and cells expressing fewer than 50 genes were removed. Additionally, we excluded cells with fewer than 300 total counts.

To ensure balanced representation of cell types and tissues, we retained the 30 most abundant cell types from the human and mouse datasets and the top 10 cell types from the fly dataset. Additionally, for the mouse and human datasets, we restricted the data to the 17 overlapping tissues shared between the two species. We also filtered out age groups with fewer than 1,000 cells to maintain statistical robustness. Only genes present in the final filtered datasets were retained.

To enable biologically meaningful comparisons across species, we mapped mouse and fly ages to human-equivalent ages. Mouse-human equivalences followed established developmental phase mappings from Flurkey, Currer, and Harrison [72], while fly-human mappings were derived by aligning lifespan curves. The specific mappings used were:

- Mouse → Human: 1m → 12.5y, 3m → 20y, 18m → 56y, 21m → 62.5y, 24m → 69y, 30m → 81y.
- Fly → Human: 5d → 5y, 30d → 40y, 50d → 68y, 70d → 90y.

After preprocessing, the final multi-species dataset contained 793,747 cells and 2,515 genes. The human subset included 520,139 cells, the mouse subset included 171,924 cells, and the fly subset included 101,684 cells.

#### Multi-species dataset (Mouse + Human)

To evaluate ACE’s ability to generalize to previously unseen cellular contexts, we constructed a multi-species dataset by combining the TMS mouse dataset [9] with the TS human dataset [23].

We began by identifying one-to-one orthologous genes between mouse and human. To ensure high-quality input data, we excluded genes expressed in fewer than 5 cells and filtered out cells with fewer than 500 detected genes or fewer than 3,000 total counts.

To maintain consistency and comparability across species, we retained only cell types shared between mouse and human with at least 100 cells in each species. Likewise, we filtered tissues to include only those represented by at least 100 cells in both species. This filtering yielded 40 common cell types and 15 shared tissues.

We selected the top 2,000 highly variable genes using the Seurat v3 method, applying batch correction via the batch key on the raw count layer. In addition to these highly variable genes, we included aging-associated genes identified by Zhang et al. [13] to augment aging-related signatures.

To enable age-aligned comparisons across species, we mapped mouse chronological ages to their human-equivalent values using established cross-species age conversion tables [72].

The resulting dataset consists of 513,924 cells and 3,991 genes across 40 cell types and 15 tissues. Among these, 15 cell types with sufficient cell numbers and age coverage were designated as held-out sets for evaluating ACE’s cross-cell-type generalization capability.

### Evaluation metrics

We used quantitative metrics implemented in the `scikit-learn` Python package to assess model performance. To enable visual comparisons across models and metrics, we generated overview tables following the format of contrastiveVI [21], where individual scores are shown as circles and aggregate scores as bars. All metrics were min-max scaled to allow cross-metric comparisons. These scaled values were then averaged into two composite scores: *Age Spatial Autocorrelation*, which quantifies the coherence of learned aging signatures, and *Cell Trait Mix Scores*, which evaluate how well cells from different background traits (e.g., cell types, tissues, sexes) are intermixed in the learned age space. A final overall score was computed by averaging these two aggregates.

For *Age Spatial Autocorrelation*, we computed both Geary’s C and Moran’s I to quantify the spatial autocorrelation of the aging variables learned by the global ACE model. While Moran’s I captures broader global trends in spatial structure, Geary’s C is more sensitive to local differences among neighboring cells.

For *Cell Trait Mix Scores*, we computed the Entropy of Mixing, which measures the diversity of background groups among the nearest neighbors of each cell in the learned age space. High entropy indicates that cells from different groups are well mixed, suggesting that the learned representation successfully removes background-specific variation. This is desirable for isolating aging signatures that generalize across different cellular contexts.

#### Geary’s C

To quantify local spatial autocorrelation in the learned aging latent space, we compute Geary’s C:

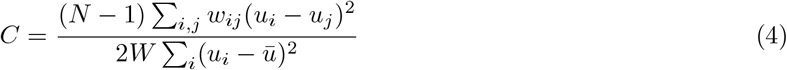

Here, *u*_*i*_ denotes the aging variable for cell *i*, ū is its global mean, *w*_*ij*_ is the spatial weight between cells *i* and *j*, and *W* = ∑_*I,j*_ *w*_*ij*_. We define *w*_*ij*_ = 1 if cell j is among the 50 nearest neighbors of cell *i*, and 0 otherwise. Since lower values of *C* indicate stronger spatial autocorrelation, we report 1 — *C* to maintain consistency with other metrics where higher values indicate better performance.

#### Moran’s I

To assess global spatial autocorrelation in the aging latent space, we compute Moran’s I:

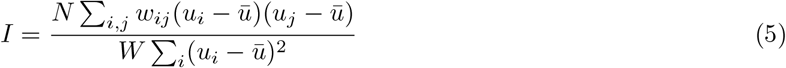

where *u*_*i*_ is the aging variable for cell *i*, ū is the global mean, and *w*_*ij*_ is defined as above. The denominator normalizes the spatial covariance between neighboring cells by the overall variance. Higher values of *I* indicate stronger spatial structure across the entire dataset.

#### Entropy of Mixing

To quantify how well cells from different background covariates (e.g., cell types, tissues, sexes) are mixed in the learned embedding space, we compute the entropy of mixing as described in prior works [15, 92]. Specifically, for each cell *<u>U</u>*, we calculate the empirical distribution of background groups among its 50 nearest neighbors, denoted by *B*_*U*_. Let p_*i*_ represent the proportion of neighbors from group *<u>i</u>* among the *c* possible groups, such that 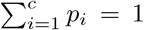. The local entropy of mixing for cell *U* is computed as:

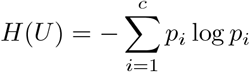

We repeat this process over 100 randomly selected cells and report the average entropy. Higher values of this metric indicate stronger mixing of background groups, reflecting better removal of unwanted variation in the embedding.

### Interpretability of the ACE Model

To interpret gene-level contributions to predicted biological age, we applied the *Expected Gradients* (EG) method [28]. Expected Gradients extends Integrated Gradients by averaging over a distribution of background samples, producing more stable and representative feature attributions for deep neural networks.

#### Definition

Given a model *f* that maps an input gene expression vector *x* ∈ ℝ^*G*^ to a scalar output (in our case, predicted biological age â), the Expected Gradient attribution for gene *g* is defined as:

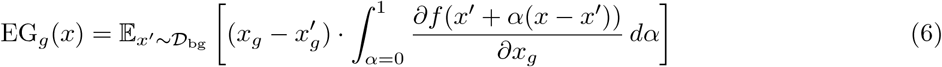

Here: - *x* is a foreground (target) sample, - *x*^′^ is drawn from a background distribution *D*_bg_, - *g* indexes the genes, - and the gradient is taken with respect to input gene *g*.

#### Foreground and Background Samples

To compute EG values in our framework: - **Foreground samples** are 200 cells randomly sampled from each tissue-cell-type pair. These represent the cells for which we wish to interpret the predicted biological age. - **Background samples** are 800 cells drawn from the same tissue-cell-type pair, representing the reference distribution *D*_bg_.

#### Application to Global and Local Models

For the *global aging model*, all cells from multiple tissue-cell-type pairs were pooled, and EG values were computed using aggregated foreground and background samples. Gene attributions were then obtained by averaging the absolute EG values across all foreground samples.

For the *local aging model*, we computed EG values separately within each tissue-cell-type pair using the respective foreground and background sets. The absolute EG values were averaged across the foreground cells in each pair. Genes were then ranked according to their mean attributions, reflecting their local importance in age prediction.

### Pathway enrichment analysis

Pathway enrichment analysis is a computational procedure used to determine whether a predefined set of genes (i.e., a biological pathway) shows statistically significant expression changes between different biological states. We used the open-source `GSEAPY` Python package for this analysis, applying the `prerank` method. Pathways with a false discovery rate (FDR) q-value below 0.05, adjusted using the Benjamini-Hochberg procedure, were considered significantly enriched and are reported in this study.

The `prerank` method is a variant of Gene Set Enrichment Analysis (GSEA) that operates on a pre-ordered list of genes instead of raw expression data. In our study, genes are ranked based on Expected Gradient values, which reflect their biological relevance. GSEA then evaluates whether genes from a particular pathway are disproportionately located near the top or bottom of the ranked list, indicating significant enrichment beyond random chance.

We conducted enrichment analysis across the following gene set libraries:

- **Aging Perturbations from GEO up/down:** Curated gene sets capturing consistent aging-associated upregulation or downregulation across multiple studies in the Gene Expression Omnibus (GEO), enabling cross-dataset evaluation of aging signatures.
- **KEGG (Kyoto Encyclopedia of Genes and Genomes):** A structured database of biological pathways encompassing metabolism, signal transduction, and disease mechanisms.
- **Reactome:** A manually curated knowledgebase detailing molecular events in signal transduction, immune function, gene regulation, and other cellular processes.
- **GO (Gene Ontology):** Hierarchical gene annotation system that organizes gene functions into three broad domains: Biological Process (BP), Cellular Component (CC), and Molecular Function (MF).

### Wet lab validation in *C. elegans*

To assess the biological relevance of aging-associated genes identified by ACE, we performed RNA interference (RNAi) knockdown experiments in *Caenorhabditis elegans* and evaluated their effects on lifespan. Specifically, we selected 8 mouse genes from among the top 150 most important global aging genes identified by the ACE global aging model: *Ppia, Ftl1, Gpx3, Rpl37, Serpina3k, Tpt1, Rps27a*, and *Uba52*. These genes were prioritized based on their relevance to proteostasis, evolutionary conservation across species, and feasibility for functional testing in *C. elegans*, avoiding those known to cause larval arrest or lethality upon knockdown.

Each mouse gene was mapped to its corresponding *C. elegans* ortholog(s), resulting in a total of 15 *C. elegans* genes tested. RNAi knockdown was carried out individually for each ortholog, and survival curves were compared to those of animals fed with an empty vector (EV) control.

### Animal Husbandry and Experimental Setup

Wild-type hermaphrodite *C. elegans* (strain N2 CGCb) were maintained under standard culture conditions at 20°C on nematode growth medium (NGM) plates seeded with *E. coli* OP50 [93]. Animals were synchronized using hypochlorite treatment and grown on OP50-seeded NGM plates until the late L4/young adult (YA) molt, approximately 50 hours post-feeding at 20°C. At this stage, animals were transferred to RNAi food on 12-well plates to initiate the aging experiments.

#### Lifespan Measurements

Lifespan assays were conducted using the WormBot imaging platform [94], which quantifies the time of death (defined as cessation of movement) for individual worms based on time-lapse video recordings. Each RNAi knockdown experiment was performed using sequence-validated RNAi clones in HT115 *E. coli*, grown on NGM plates supplemented with 5-fluoro-2’-deoxyuridine (FUDR) to inhibit progeny production. The FUDR and RNAi clones may also inhibit mitotic development, so worms were plated at the L4/YA stage to ensure that any observed lifespan changes were attributable to gene knockdown effects during aging rather than development. Each well contained 20-30 animals.

## Footnotes

1 https://github.com/suinleelab/ACE

