## Supplementary Appendix for "An Explainable AI Framework for Identifying Universal Aging Signatures in Cell Embeddings"

964

### Supplementary Appendix

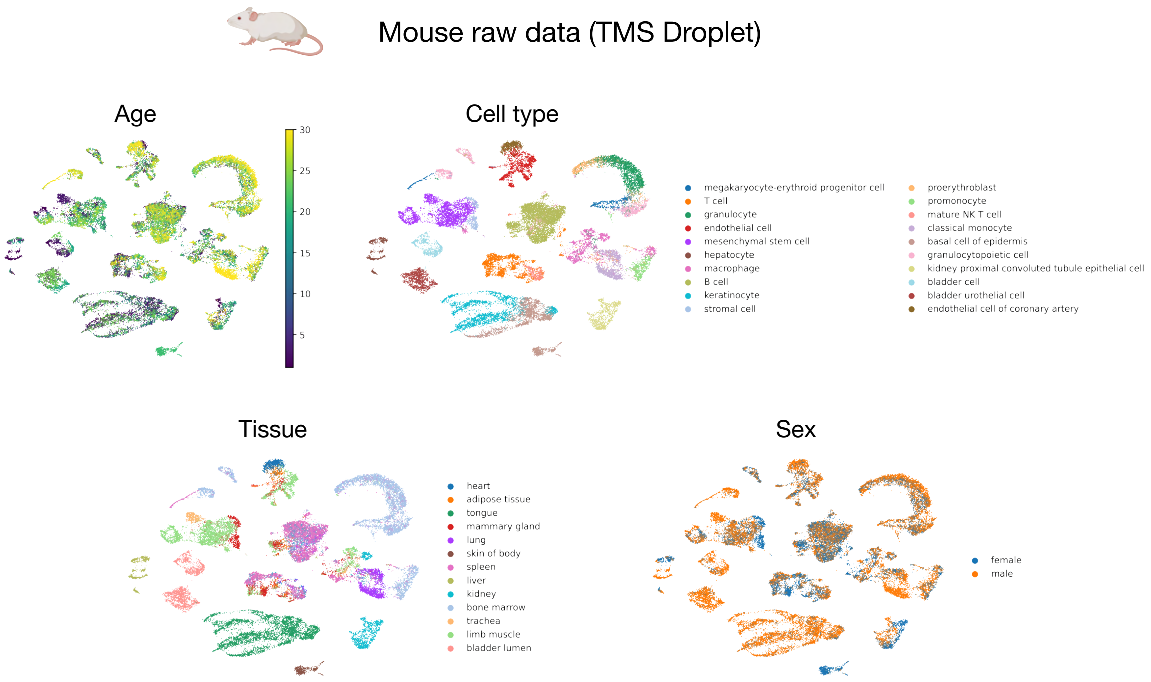


Supplementary Figure 1: **Visualization of the TMS Droplet dataset using UMAP applied to normalized count data.** Plots are colored by age (top left), cell type (top right), tissue (bottom left), and sex (bottom right).

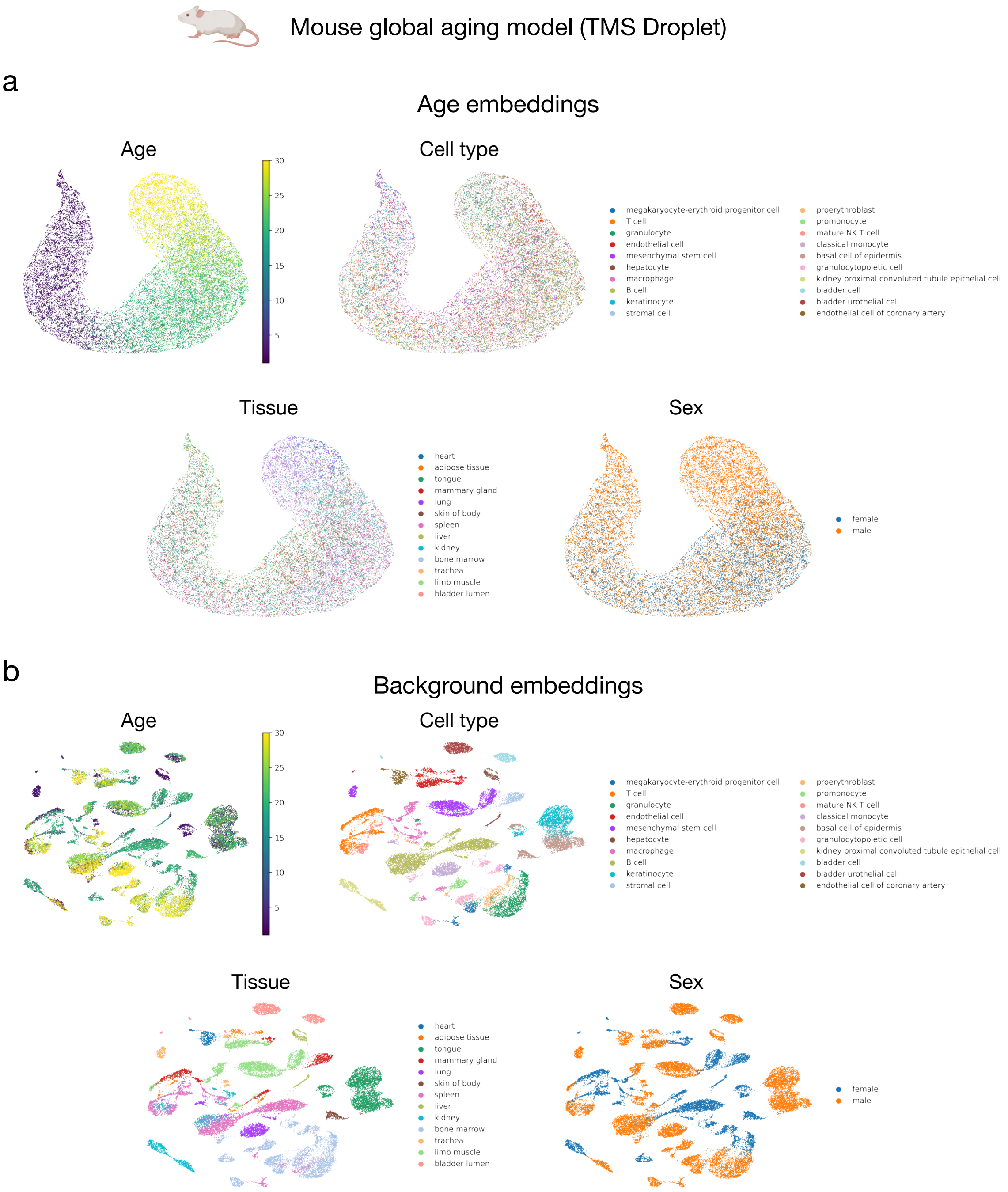


Supplementary Figure 2: **Visualization of age and background embeddings from the global aging model learned by ACE using the TMS Droplet dataset. a.** UMAP of age embeddings colored by age, cell type, tissue, and sex. **b.** UMAP of background embeddings colored by the same attributes.

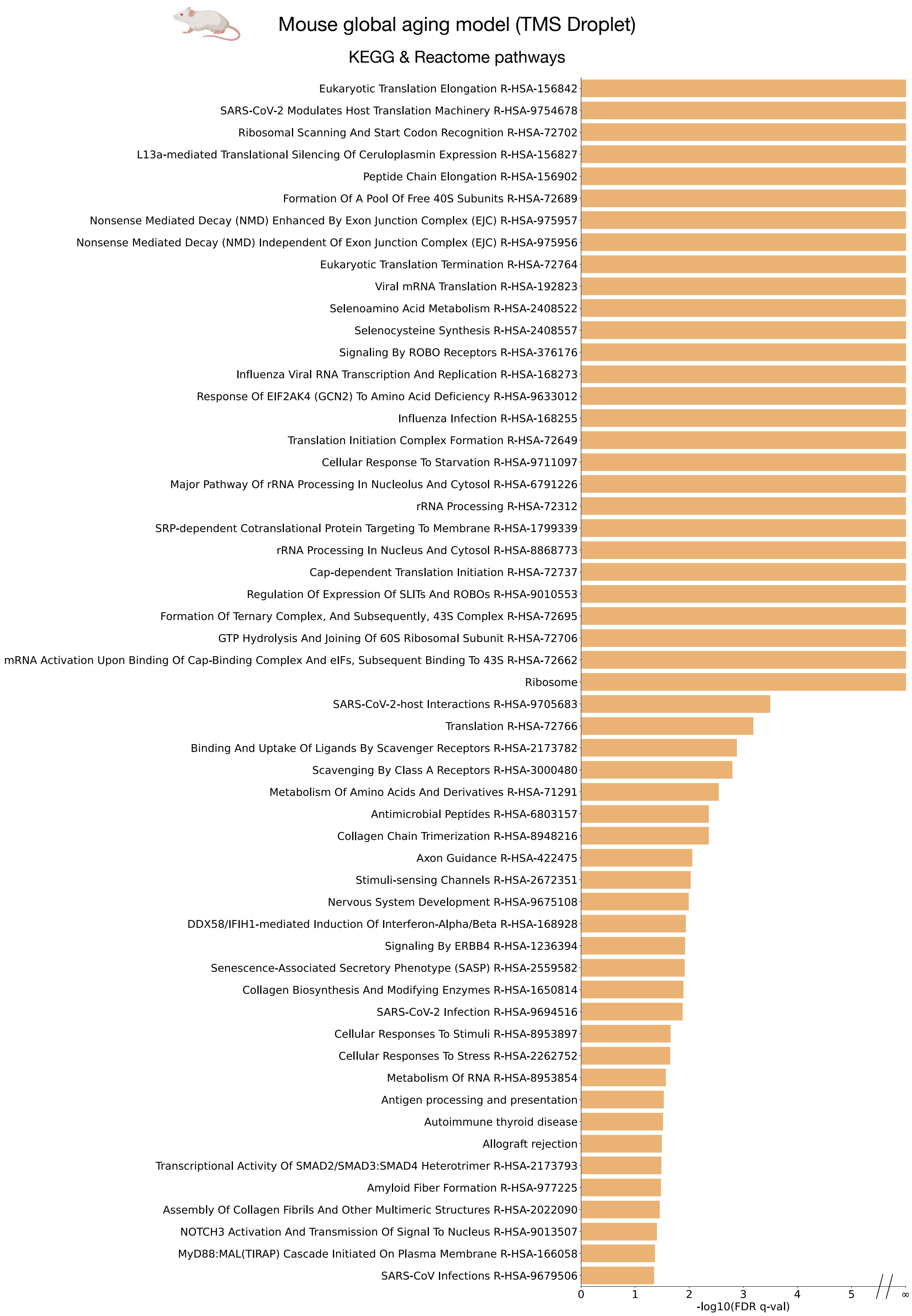


Supplementary Figure 3: **Full list of KEGG and Reactome pathways significantly enriched by the mouse global aging model on the TMS Droplet dataset.** Significance was assessed at FDR $q < 0.05$ using the Benjamini-Hochberg correction.

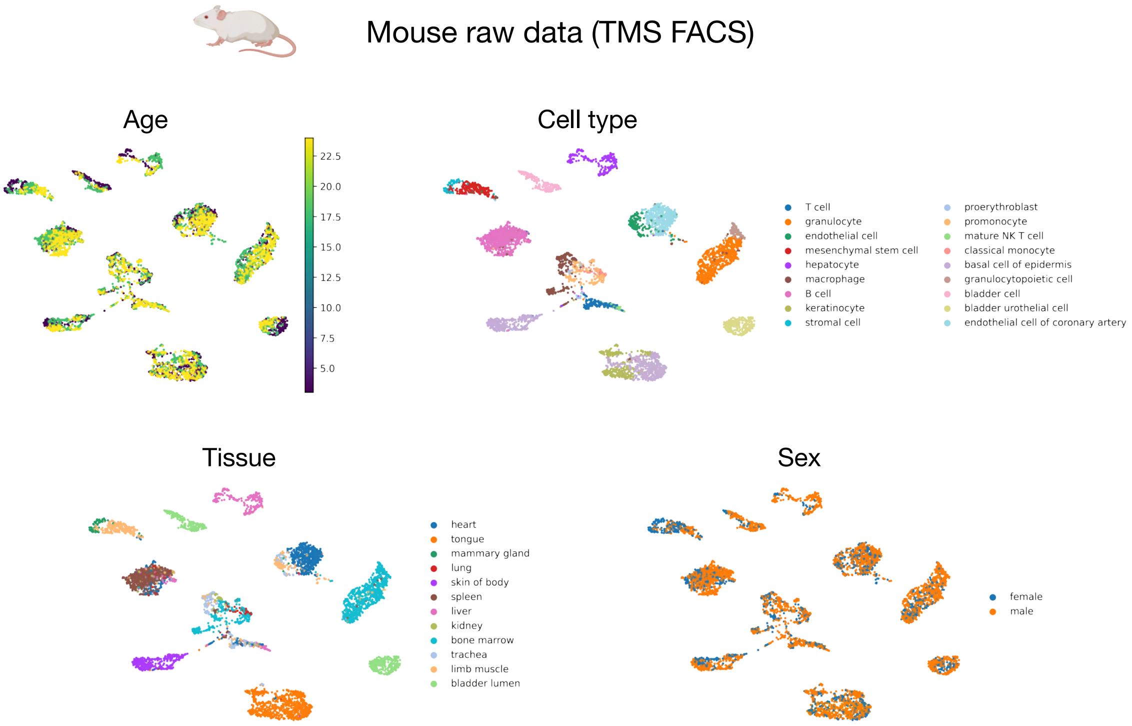


Supplementary Figure 4: **Visualization of the TMS FACS dataset using UMAP applied to normalized count data.** Plots are colored by age (top left), cell type (top right), tissue (bottom left), and sex (bottom right).

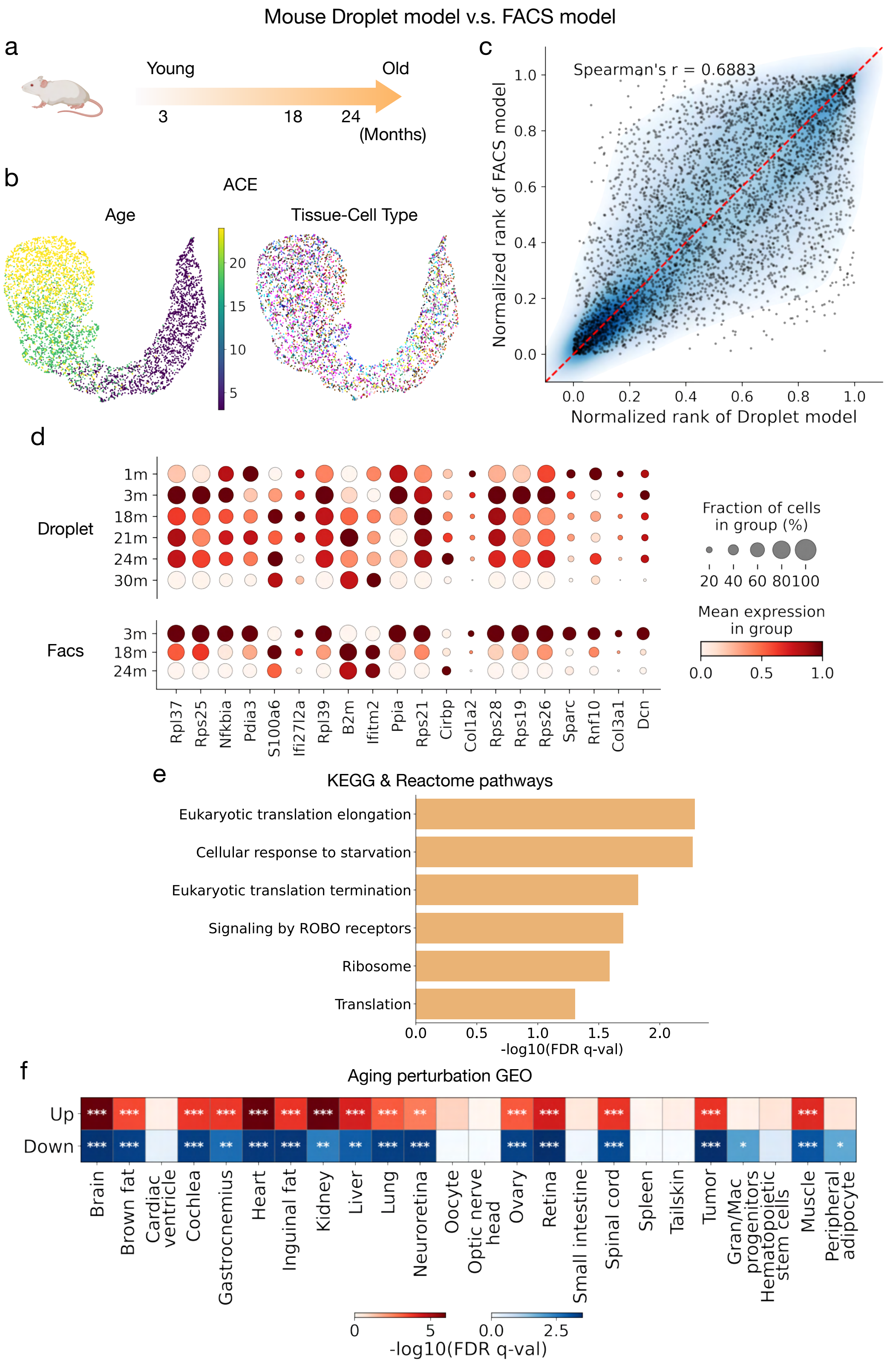


Supplementary Figure 5

Supplementary Figure 5: **Comparison of mouse Droplet and FACS models to validate global aging signatures.** **a.** Aging time points represented in the TMS FACS dataset, spanning from 3 to 24 months. The FACS dataset was processed using consistent filtering steps with the Droplet dataset, including the removal of low-quality cells, rare tissues/cell types, and donors with insufficient representation, resulting in 24,546 cells and 4,120 genes across three age groups (3m, 18m, and 24m). **b.** UMAP visualization of age embeddings learned by ACE using the TMS FACS dataset, showing that the learned embeddings capture a strong age-related gradient (left) while being disentangled from tissue and cell-type identity (right). **c.** Spearman correlation between the normalized gene rankings of the Droplet and FACS models (Spearman's $r = 0.6883$), demonstrating strong concordance and suggesting that the Droplet-derived global aging signatures are robust and reproducible in an independent dataset. **d.** Expression dynamics of selected overlapping genes from the top 150 ranked genes in both Droplet and FACS models. These genes exhibit consistent age-associated changes, with similar patterns of upregulation or downregulation across matched time points, reinforcing their roles as core aging-related genes. The size of each dot represents the fraction of cells in a group expressing the gene, while color intensity reflects the mean expression within that group. **e.** Selected overlapping KEGG and Reactome pathways identified by both models, including translation-related processes (e.g., eukaryotic translation elongation and termination), cellular response to starvation, and ROBO signaling. These representative pathways correspond to those in Figure 2h and highlight biological processes consistently implicated in aging across datasets. **f.** Gene set enrichment analysis results from the FACS model, using the "Aging Perturbations from GEO Up" and "Aging Perturbations from GEO Down" collections. These databases aggregate results from GEO studies comparing aged versus young tissues and cell types. The FACS-derived global aging gene rankings show significant enrichment in many of these curated gene sets. Red boxes indicate enrichment in *upregulated* gene sets with age, whereas blue boxes indicate enrichment in *downregulated* gene sets. Color intensity represents $-\log_{10}$(FDR $q$-value), and significance was assessed using Benjamini-Hochberg correction. Asterisks indicate significance thresholds ($^{*}p < 0.05$, $^{**}p < 0.01$, $^{***}p < 0.001$). Notably, genes upregulated with aging are strongly enriched in multiple tissues (e.g., brain, heart, muscle, adipose), while downregulated genes are observed in select tissues, confirming that the FACS model captures both shared and tissue-specific aging perturbations. Gran/Mac progenitors represent granulocyte/macrophage progenitors.

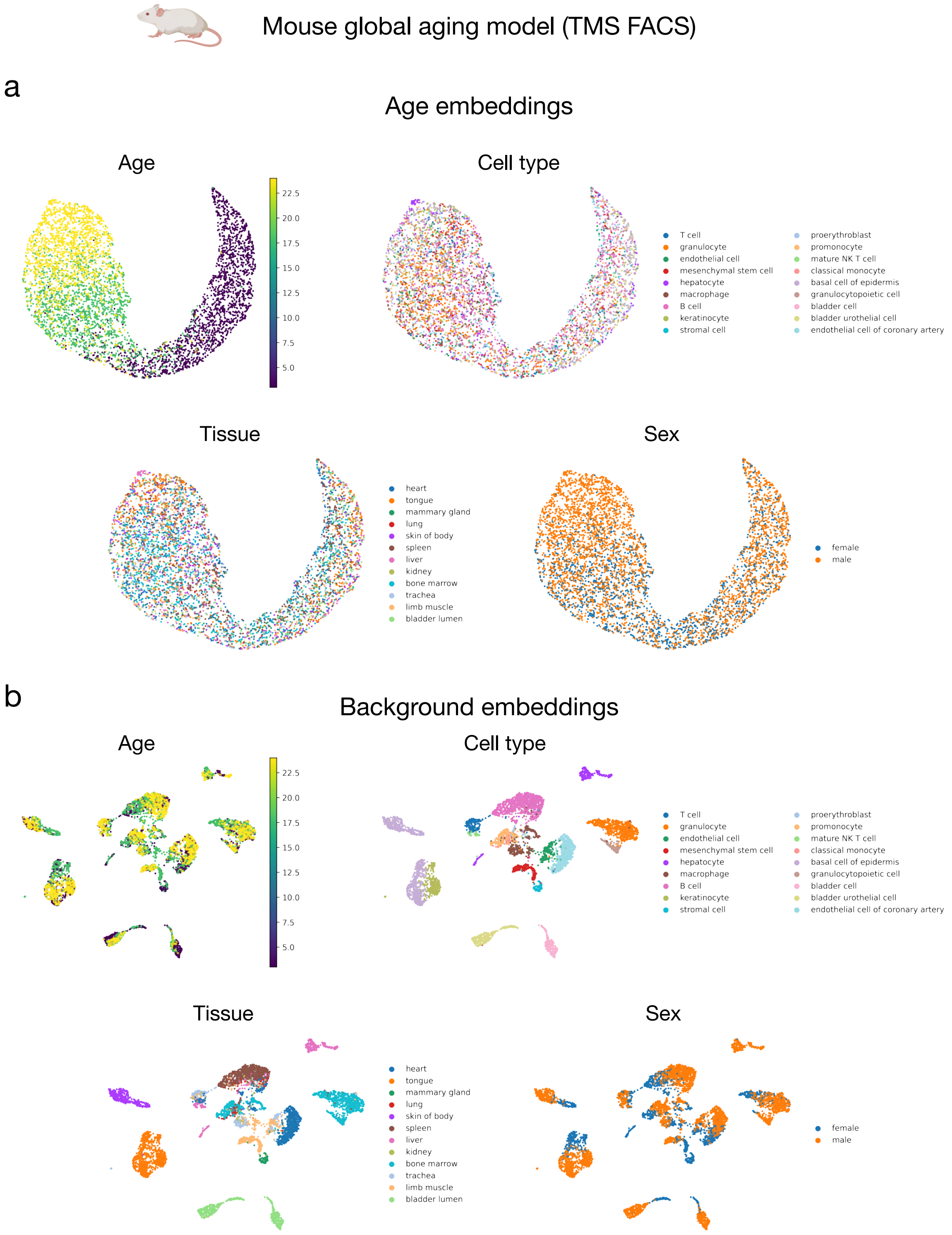


Supplementary Figure 6: **Visualization of age and background embeddings from the global aging model learned by ACE using the TMS FACS dataset. a.** UMAP of age embeddings colored by age, cell type, tissue, and sex. **b.** UMAP of background embeddings colored by the same attributes.

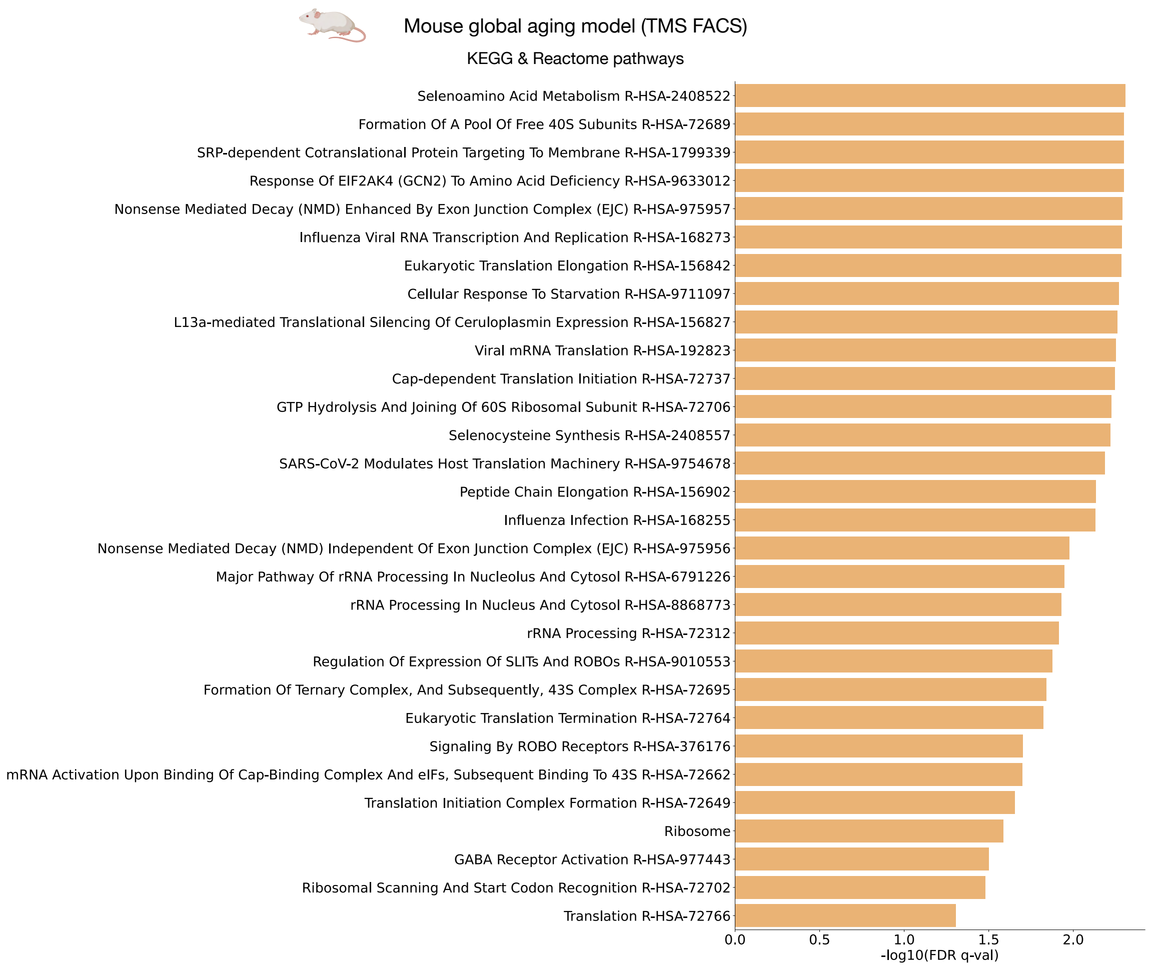


Supplementary Figure 7: **Full list of KEGG and Reactome pathways significantly enriched by the mouse global aging model on the TMS FACS dataset.** Significance was assessed at FDR $q < 0.05$ using the Benjamini-Hochberg correction. A total of 29 pathways identified in the FACS model overlapped with those discovered by the Droplet model (Supplementary Figure 3), highlighting robust and reproducible aging-associated biological processes across datasets.

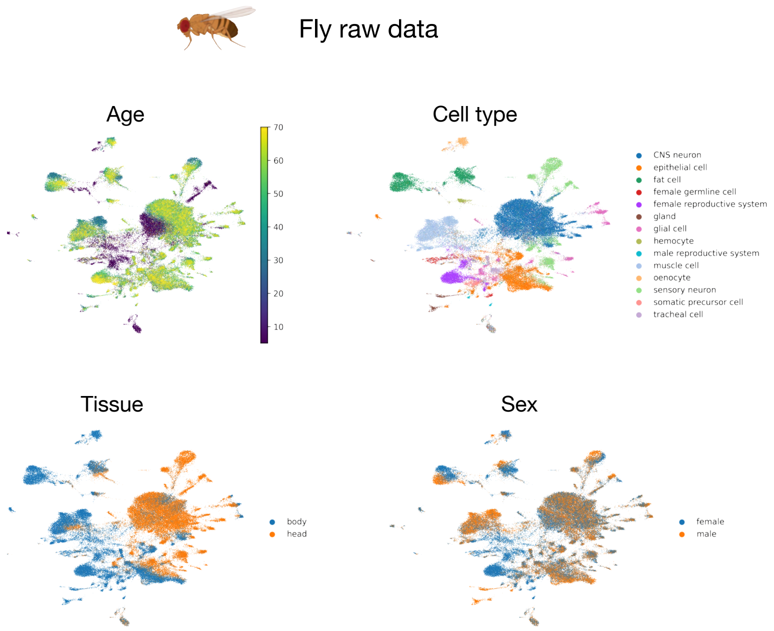


Supplementary Figure 8: **Visualization of the Aging Fly Cell Atlas using UMAP applied to normalized count data.** Plots are colored by age (top left), cell type (top right), tissue (bottom left), and sex (bottom right).

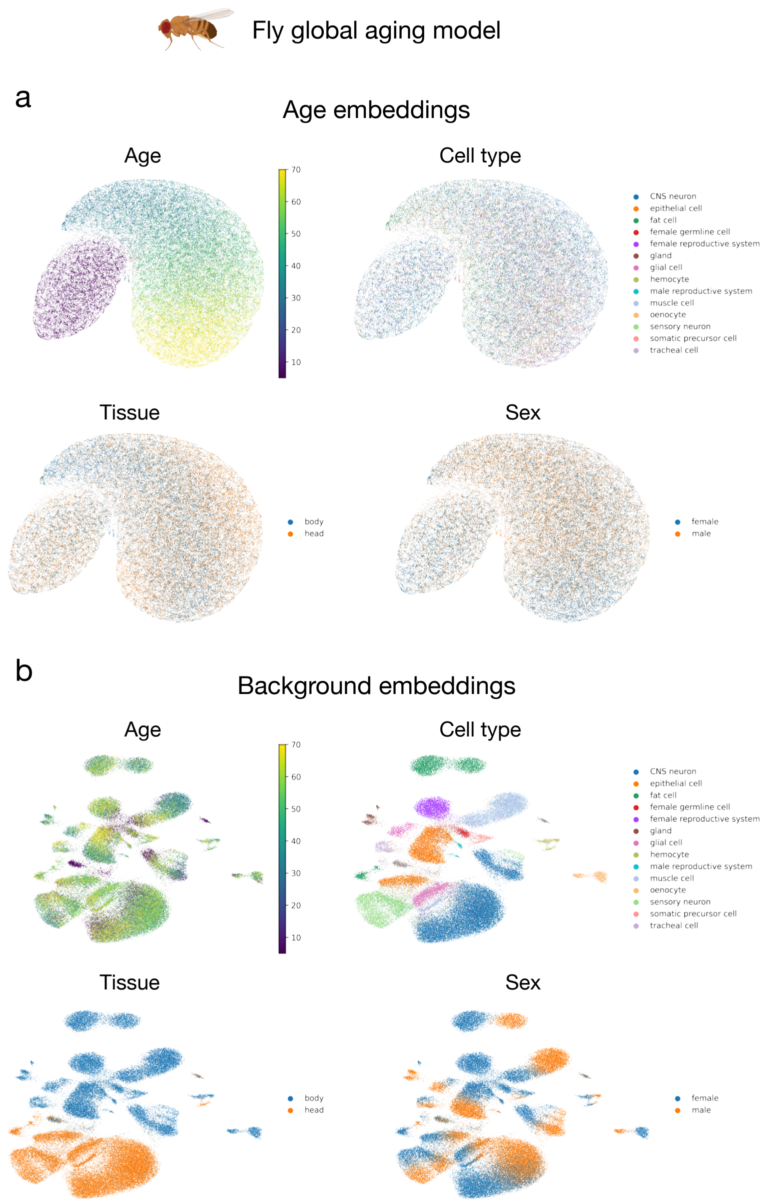


Supplementary Figure 9: **Visualization of age and background embeddings from the global aging model learned by ACE using the Aging Fly Cell Atlas. a.** UMAP of age embeddings colored by age, cell type, tissue, and sex. **b.** UMAP of background embeddings colored by the same attributes.

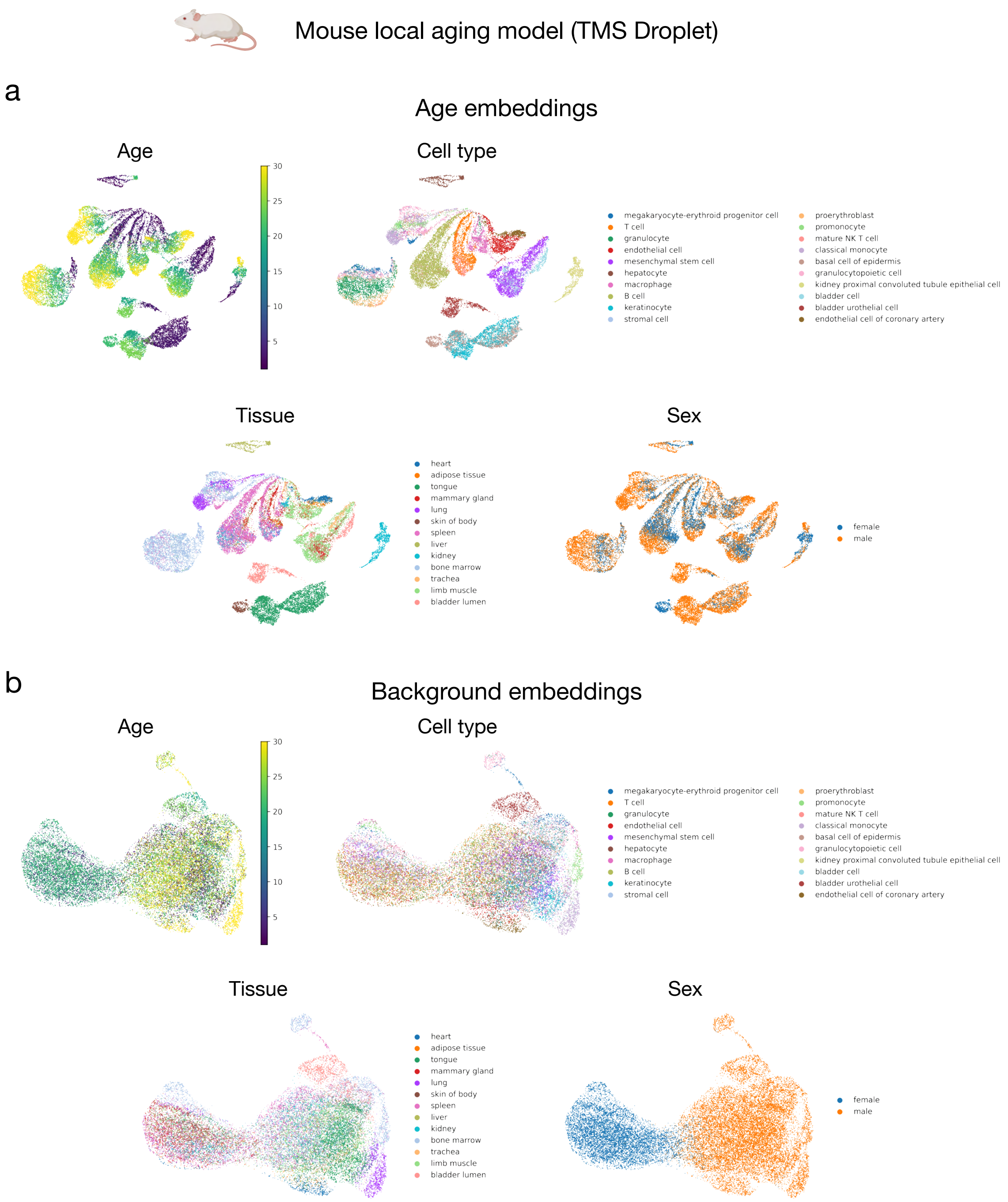


Supplementary Figure 10: **Visualization of age and background embeddings from the local aging model learned by ACE using the TMS Droplet dataset. a.** UMAP of age embeddings colored by age, cell type, tissue, and sex. **b.** UMAP of background embeddings colored by the same attributes.

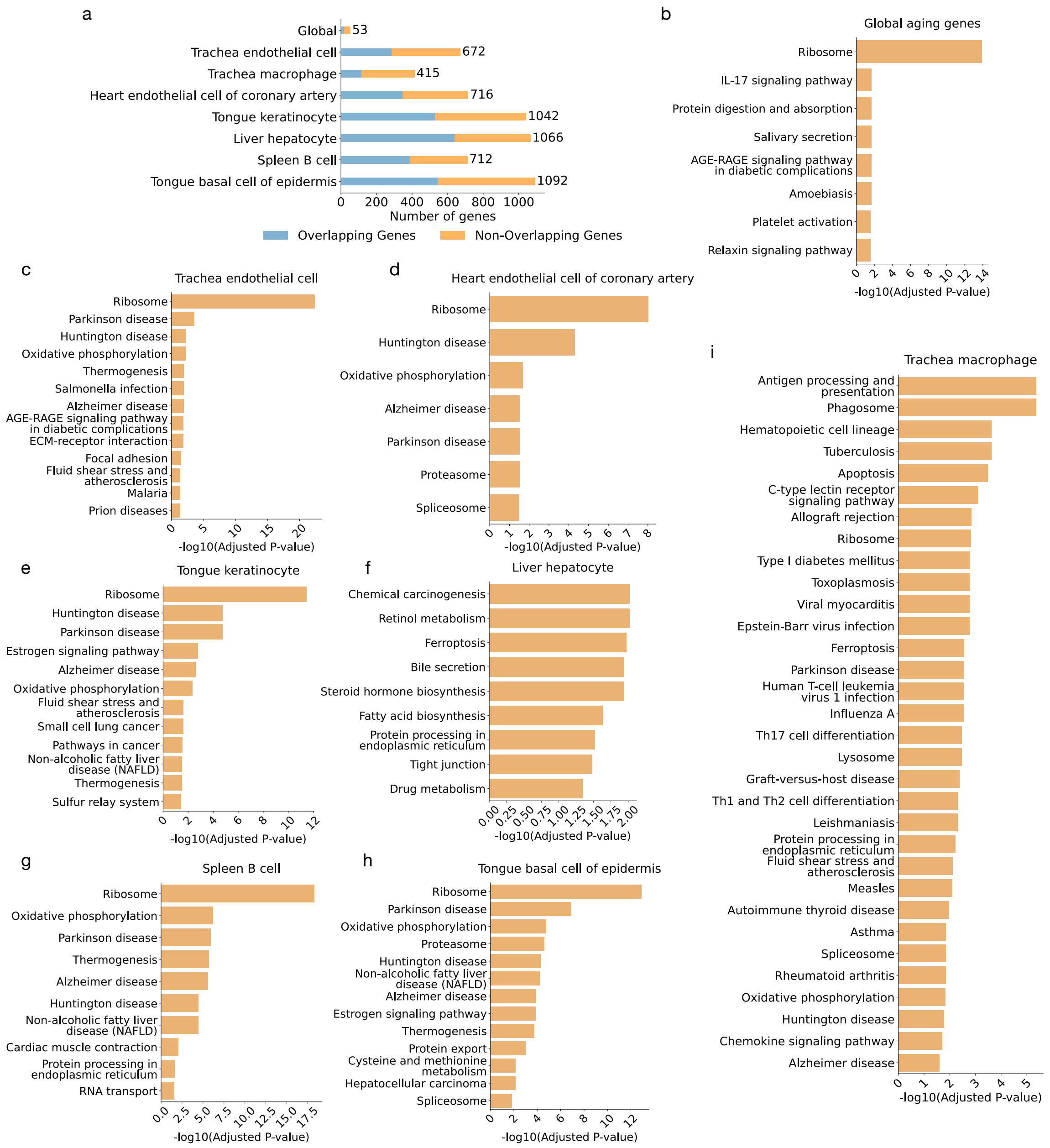


Supplementary Figure 11

Supplementary Figure 11: **Comparison of ACE-derived global and tissue-cell-type-specific aging genes with those identified by the linear model from Zhang et al. [1]. a.** Bar plot showing the number of overlapping and non-overlapping aging genes identified by the ACE model and the linear model. For each group, we matched the number of top-ranked ACE genes to the number of aging genes reported by the linear model, allowing for a direct and fair comparison between the two approaches. This comparison shows that ACE identifies a substantial number of additional aging-related genes beyond those captured by the linear model, demonstrating its improved sensitivity for detecting comprehensive aging signatures. **b-i.** Gene set enrichment analysis of KEGG pathways performed on the non-overlapping ACE genes for the global group and each tissue-cell-type-specific group. The enriched pathways are strongly associated with known hallmarks of aging. These analyses demonstrate that ACE not only captures key aging pathways identified by the linear model but also uncovers additional aging signatures missed by the linear approach. By identifying both shared global aging pathways and tissue-cell-type-specific pathways, ACE provides a deeper understanding of the complex and heterogeneous nature of the aging process. Significance was assessed at FDR $q < 0.05$ using the Benjamini-Hochberg correction.

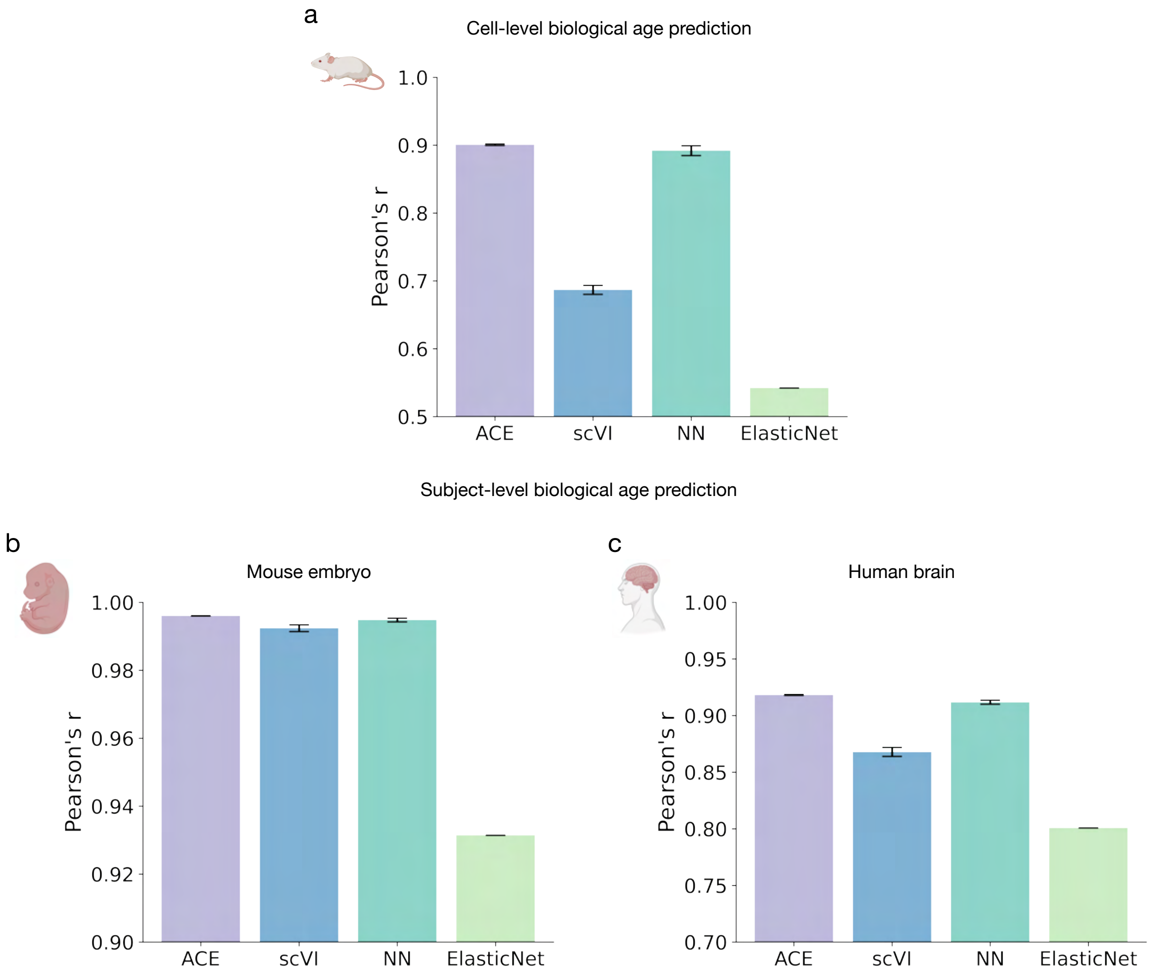


Supplementary Figure 12: **Biological age prediction performance comparison.** (a) Cell-level biological age prediction across methods. ACE achieves high accuracy compared to baseline models, including scVI, NN, and ElasticNet. (b, c) Subject-level biological age prediction for mouse embryo (b) and human brain (c) datasets. ACE consistently performs competitively or better than other methods, demonstrating its effectiveness in capturing aging-related variation at both the cell and subject levels. Error bars represent standard errors of Pearson's correlation coefficients calculated across 10 replicates.

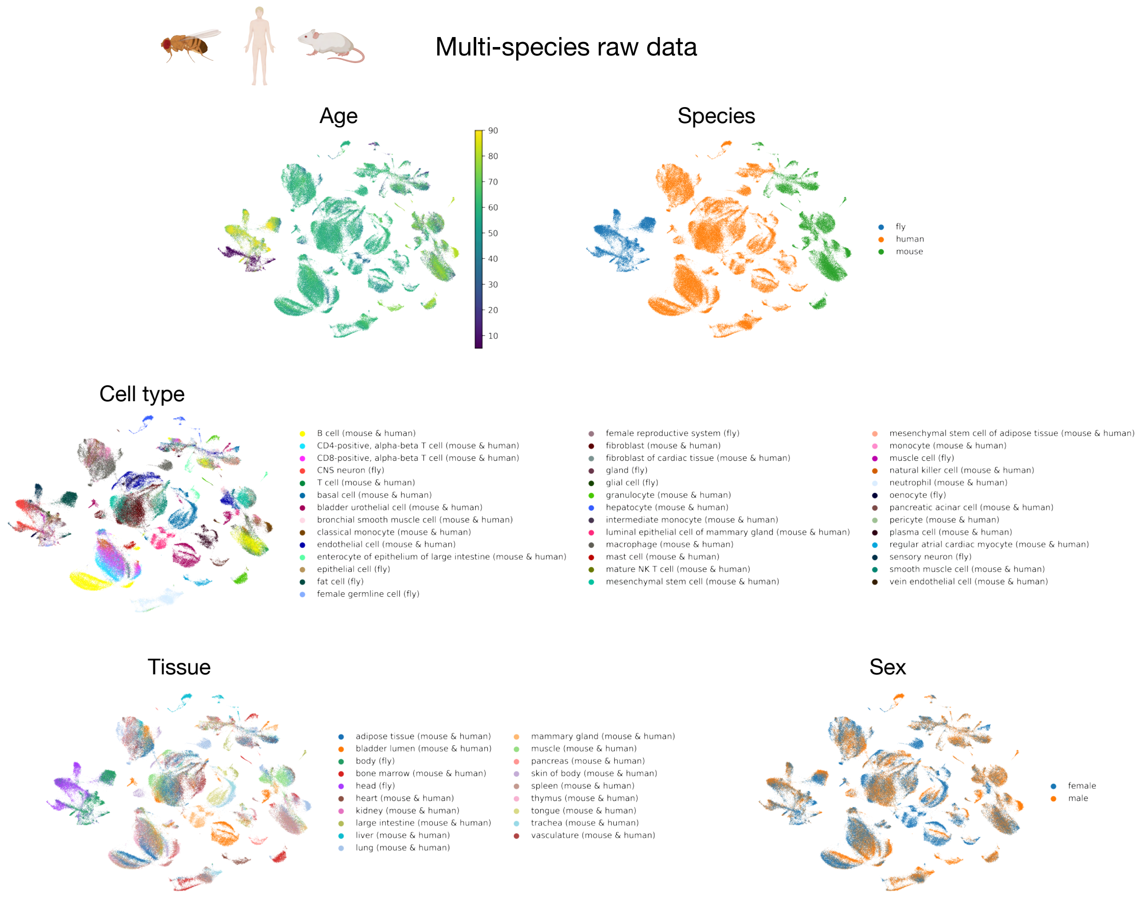


Supplementary Figure 13: **Visualization of the multi-species dataset using UMAP applied to normalized count data.** Plots are colored by age (top left), species (top right), cell type (middle), tissue (bottom left), and sex (bottom right).

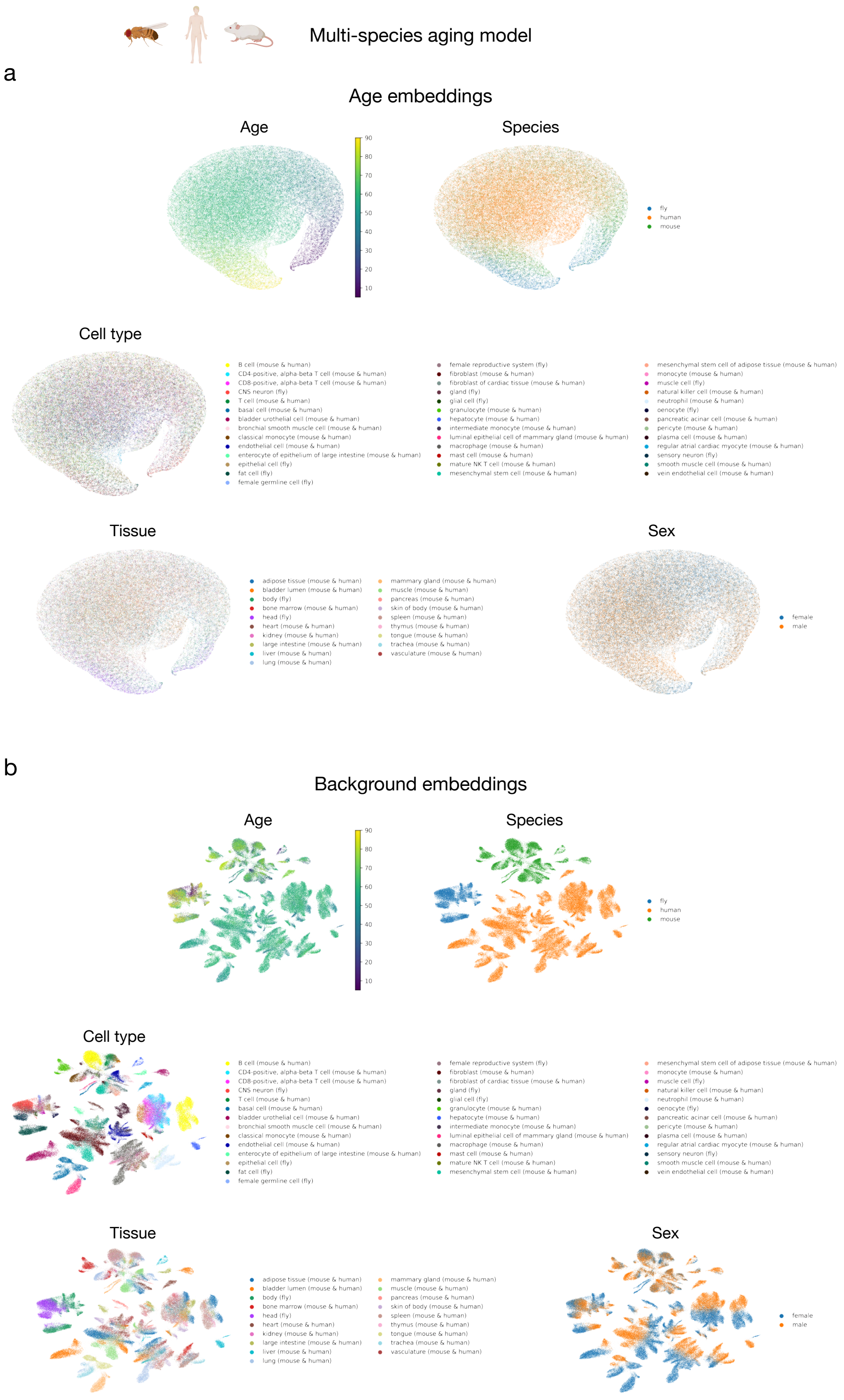


Supplementary Figure 14: **Visualization of age and background embeddings from the multi-species aging model learned by ACE. a.** UMAP of age embeddings colored by age, species, cell type, tissue, and sex. **b.** UMAP of background embeddings colored by the same attributes.

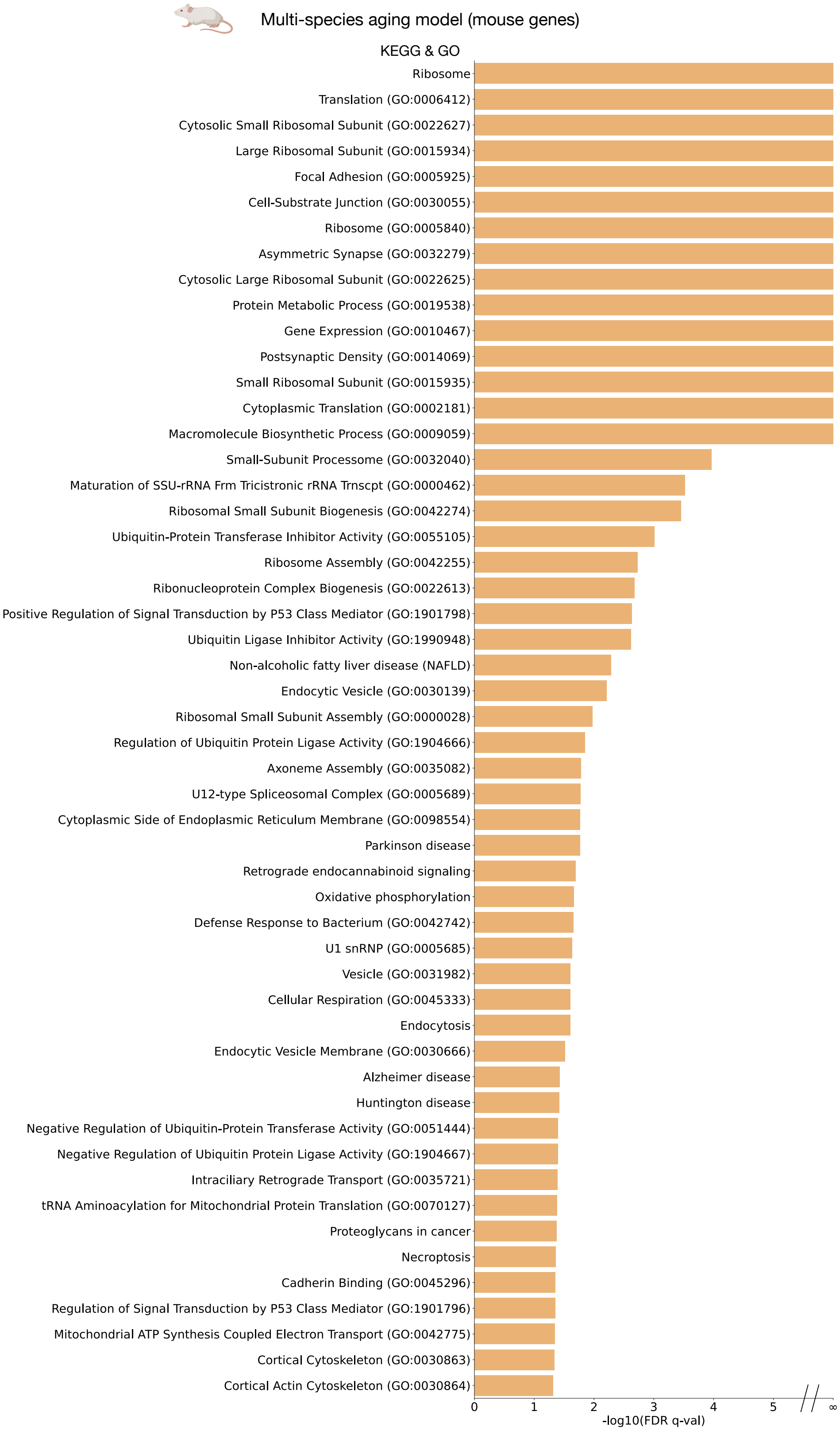


Supplementary Figure 15: **Full list of KEGG and GO pathways significantly enriched by the multi-species aging model using mouse gene sets.** Significance was assessed at FDR $q < 0.05$ using the Benjamini-Hochberg correction.

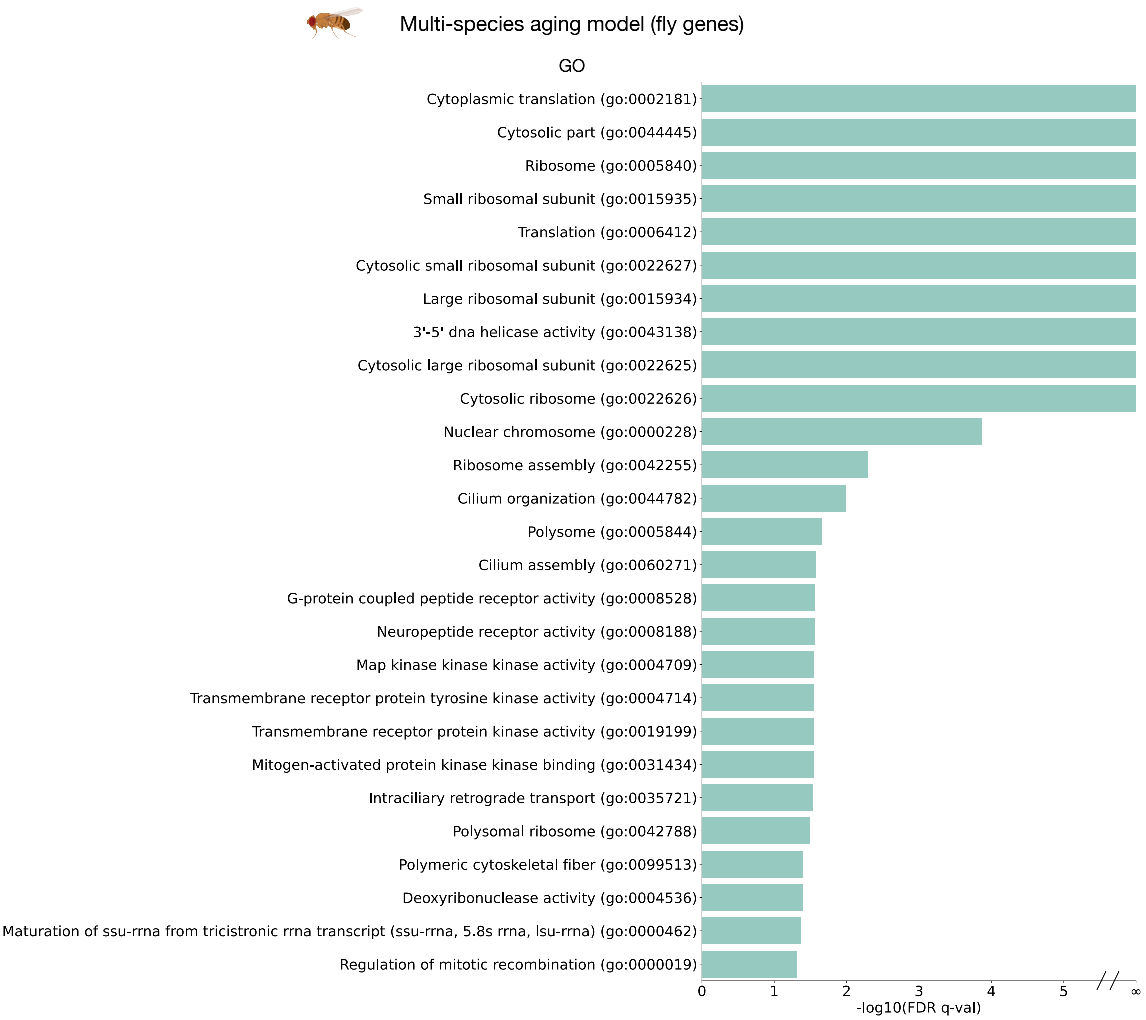


Supplementary Figure 16: **Full list of GO pathways significantly enriched by the multi-species aging model using fly gene sets.** Significance was assessed at FDR $q < 0.05$ using the Benjamini-Hochberg correction.

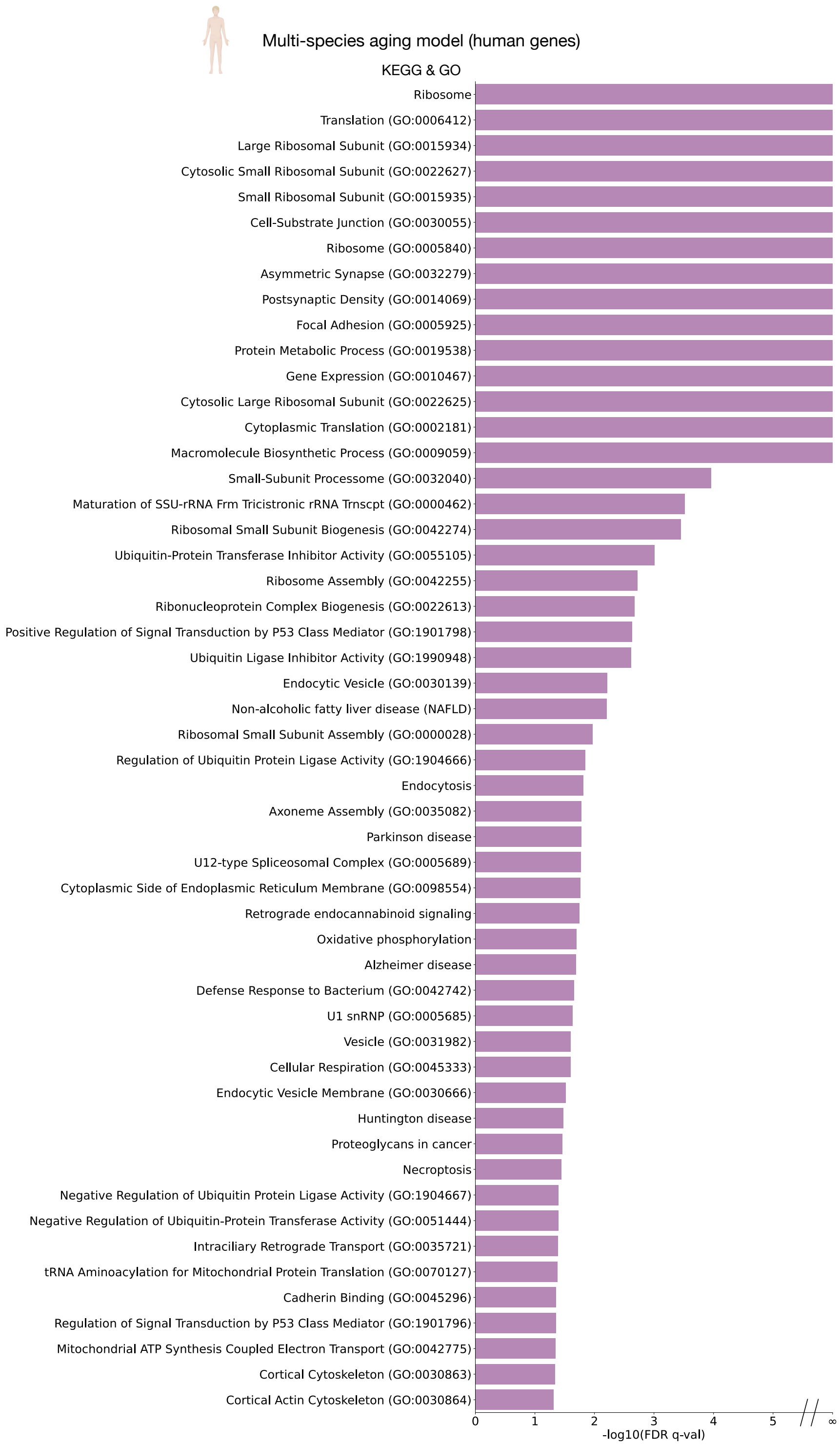


Supplementary Figure 17: **Full list of KEGG and GO pathways significantly enriched by the multi-species aging model using human gene sets.** Significance was assessed at FDR $q < 0.05$ using the Benjamini-Hochberg correction.

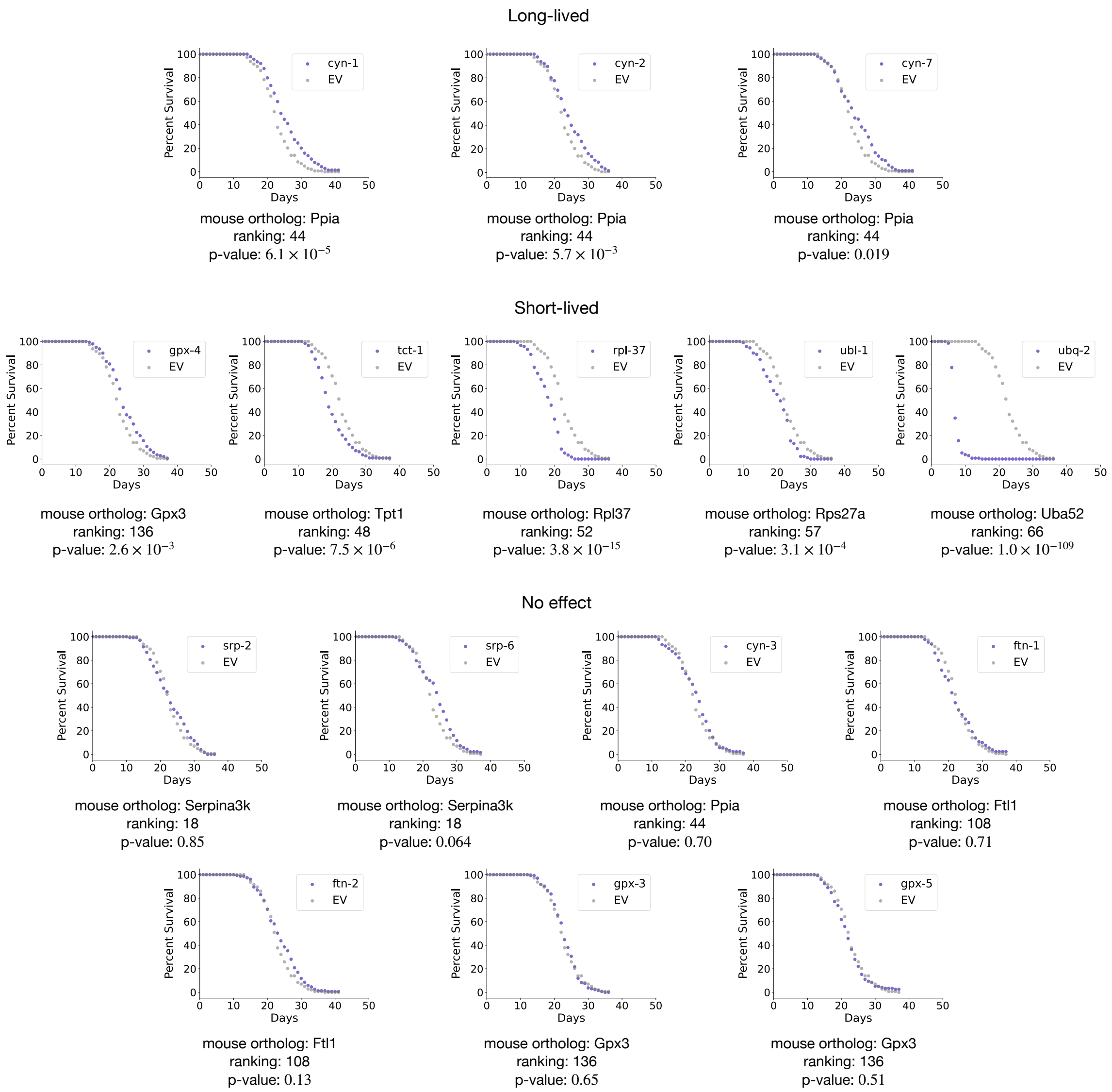


Supplementary Figure 18: **Wet lab validation of ACE-identified aging genes using C. elegans RNAi lifespan assays.** Survival curves for knockdown of *C. elegans* orthologs corresponding to selected global aging genes from the mouse global ACE model. Each plot shows the percent survival of animals treated with gene-specific RNAi (purple) compared to empty vector (EV) control (gray). Mouse gene names, their rankings from the ACE global aging model, and p-values from t-tests (adjusted using the Benjamini-Hochberg correction) are displayed below each plot. Genes are grouped into three categories based on their effects on lifespan: **Long-lived** (knockdown extends lifespan), **Short-lived** (knockdown shortens lifespan), and **No effect** (no significant impact on lifespan).
